# Global discovery of bacterial RNA-binding proteins by RNase-sensitive gradient profiles reports a new FinO domain protein

**DOI:** 10.1101/2020.06.22.164913

**Authors:** Milan Gerovac, Youssef El Mouali, Jochen Kuper, Caroline Kisker, Lars Barquist, Jörg Vogel

## Abstract

RNA-binding proteins (RBPs) play important roles in bacterial gene expression and physiology but their true number and functional scope remain little understood even in model microbes. To advance global RBP discovery in bacteria, we here establish glycerol gradient sedimentation with RNase treatment and mass spectrometry (GradR). Applied to *Salmonella enterica*, GradR confirms many known RBPs such as CsrA, Hfq and ProQ by their RNase-sensitive sedimentation profiles, and discovers the FopA protein as a new member of the emerging family of FinO/ProQ-like RBPs. FopA, encoded on resistance plasmid pCol1B9, primarily targets a small RNA associated with plasmid replication. The target suite of FopA dramatically differs from the related global RBP ProQ, revealing context-dependent selective RNA recognition by FinO-domain RBPs. Numerous other unexpected RNase-induced changes in gradient profiles suggest that cellular RNA helps to organize macromolecular complexes in bacteria. By enabling poly(A)-independent generic RBP discovery, GradR provides an important element in the quest to build a comprehensive catalogue of microbial RBPs.

## INTRODUCTION

As they do in the other kingdoms of life, RNA-binding proteins (RBPs) play vital roles in many cellular processes in bacteria. Their activities range from enabling basic protein synthesis to facilitating regulatory RNA networks required for bacterial adaptation and genome defence (Holmqvist and Vogel 2018; Babitzke et al. 2019). However, while the collective results of recent global screens have produced nearly saturated RPB catalogues in several model eukaryotes (Hentze et al. 2018), our knowledge about bacterial RBPs has remained patchy, having accumulated largely through studies of individual proteins and serendipitous discoveries. As a result, *Escherichia coli* has ~180 annotated RBPs (Holmqvist and Vogel 2018), and far fewer are known for other bacterial species. At the same time, the surging interest in microbiomes keeps increasing the number of bacteria with relevance to human health (Browne et al. 2016; Lagier et al. 2016). An understanding of RNA-centric regulation and the importance of RNA-protein interactions in these many understudied species demands new global approaches to identify bacterial RBPs in a generic manner.

Experimental screens for bacterial RBPs have been hampered by the lack of two important features that have accelerated global RBP discovery in eukaryotes: functional poly(A) tails on transcripts, and efficient incorporation of crosslink-enhancing artificial nucleotides (Bao et al. 2018; Hör et al. 2018). However, there has recently been a surge in poly(A)-independent RBP discovery methods (Smirnov et al. 2016; Asencio et al. 2018; Queiroz et al. 2019; Shchepachev et al. 2019; Trendel et al. 2019; Urdaneta et al. 2019), an example of which is Grad-seq, which predicts new RNA-protein complexes by RNA-seq and mass spectrometry (MS) of cellular lysates after fractionation on glycerol gradients (Smirnov et al. 2016). Applied to *Salmonella enterica* serovar Typhimurium (henceforth *Salmonella*), Grad-seq identified the protein ProQ as the third major RBP to be associated with small regulatory RNAs (sRNAs) (Smirnov et al. 2016), after the Sm-like protein Hfq and the translational repressor CsrA.

ProQ belongs to an emerging class of RBPs whose hallmark is a conserved FinO domain (G. Chaulk et al. 2011; Attaiech et al. 2016; Attaiech et al. 2017; Olejniczak and Storz 2017; Holmqvist and Vogel 2018; Babitzke et al. 2019). This RBP is exciting as it binds hundreds of different *E. coli* and *Salmonella* mRNAs and sRNAs with no obvious consensus sequence (Smirnov et al. 2016; Holmqvist et al. 2018; Melamed et al. 2020), which raises the possibility of FinO domain-mediated global post-transcriptional control via the recognition of a complex structural RNA code (Gonzalez et al. 2017; Holmqvist et al. 2018).

ProQ ranked high in several recent proof-of-concept experiments applying eukaryotic poly(A)-independent RBP enrichment protocols to enteric bacteria. These protocols generally used organic extraction steps or silica-based solid-phase purification to specifically recover RBPs after *in vivo* UV-crosslinking (Asencio et al. 2018; Queiroz et al. 2019; Shchepachev et al. 2019; Trendel et al. 2019; Urdaneta et al. 2019). Notwithstanding the great potential of these methods, a quick cross-comparison of the published results suggests that each of them has their own bias and dropout rate (Smith et al. 2020). For example, instead of getting enriched, CsrA was depleted upon UV crosslinking in some of these studies (Queiroz et al. 2019; Urdaneta et al. 2019). Thus, additional approaches are needed to unravel the full scope of RBPs in bacteria.

Here, we present such an alternative approach termed GradR, which predicts bacterial RBPs through their changed sedimentation profile in a glycerol gradient when associated RNA partners are removed. We demonstrate proof-of-principle by unveiling the YafB protein of previously unknown function as the third FinO-domain RBP in *Salmonella*. Like FinO, the founding member of this RBP family (Biesen and Frost 1994), YafB is encoded on a plasmid, and we have renamed it here FopA (<u>F</u>in<u>O</u> domain protein on plasmid/phage <u>A</u>). Intriguingly, the target transcripts of FopA are distinct from both, those of FinO and ProQ, suggesting that the highly conserved FinO domain recognizes very different RNAs despite the fact that all these proteins act in the same cytosolic environment. Furthermore, RNase-treatment causes other intriguing changes in in-gradient distributions, indicating a much more prominent role of RNA in organizing the cellular proteome of bacteria than currently appreciated.

## RESULTS

### Differential sedimentation profiles of RBPs upon prior RNase digestion

GradR builds on control experiments for Grad-seq (Smirnov et al. 2016), which indicated that if cellular extracts were pre-treated with ribonuclease (RNase), this would free proteins of RNA ligands and, in a glycerol gradient, shift RBPs to lighter fractions (Fig. 1A). RBPs will then be identified by comparative mass spectrometry of the treated and untreated gradients, so the theory. As do the conceptually related R-DeeP (Caudron-Herger et al. 2019) and DIF-FRAC (Mallam et al. 2019) approaches for RBP discovery in eukaryotes, GradR works without a UV-crosslinking step.

**Fig. 1.**
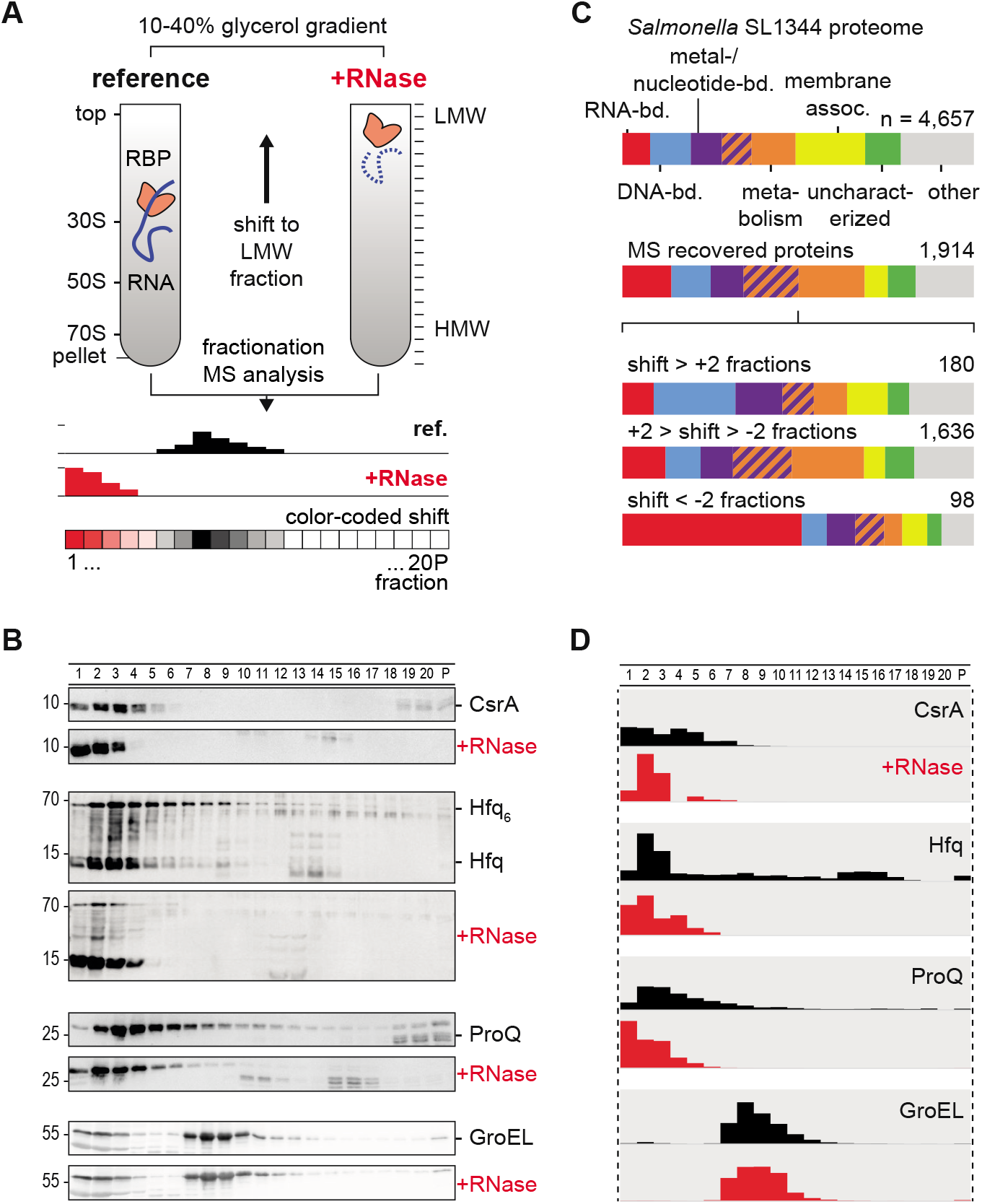
RNase-dependent gradient fraction reveals RNA-binding character of proteins. (A) The *Salmonella* lysate was digested with RNase A/T1, and sedimented in a glycerol gradient that yielded RNA-dependent shifts for RNA-binding proteins (RBPs). The profile shows changes in protein abundance, color-coded from black (shifting from) to red (shifting towards). (B) Sedimentation profile of the global RNA-binding proteins CsrA, Hfq, and ProQ were detected by FLAG-tagged variants and Western blotting. After RNase treatment, RBPs shifted to top fractions. The Hfq monomer accumulated after RNase digestion in top fractions. GroEL sedimentation was not affected by RNase treatment. n=2 (C) One third of the *Salmonella* proteome was recovered by MS. RNA-binding proteins (red) were enriched in the group of proteins with a high relative shift to the top (< −2 fractions). n=2 (D) The global RBPs correlated well in MS quantification with Fig. 1b. Protein abundance was normalized to total protein levels per gradient. Reference and RNase treated sedimentation profiles were represented as black and red bars, respectively. n=2

We first optimized the treatment of bacterial lysates for rapid degradation of RNA while keeping proteolysis to a minimum, settling at nuclease concentrations of 0.3 μg/μl RNase A and 0.8 U/μl RNase T1, and a 20-min reaction at 20 °C for a *Salmonella* lysate of an A_260_ of ~150. Gradients were prepared with 10 mM magnesium concentration in order to ensure 70S ribosome integrity and mRNA co-migration with 70S ribosomes or polysomes (Gros et al. 1961; Zitomer and Flaks 1972) in the reference sample. Probing for the model RBPs CsrA, Hfq, and ProQ on western blots, we observed the expected RNase-induced shifts towards low-molecular weight (LMW) fractions (Fig. 1B). In the case of Hfq, the RNase treatment converted most of the hexameric form to monomers that accumulated in early LMW fractions, supporting previous suggestions (Argaman et al. 2012; Panja and Woodson 2012) that of Hfq. As expected, the protein chaperone GroEL serving as a negative control did not show an RNase-induced shift.

### Global analysis of RNase-treated gradient fractions

For a global RBP prediction, we determined protein sedimentation profiles by mass spectrometry (MS) across the 20 gradient fractions plus the pellet and compared these profiles between the RNase-treated and untreated reference samples. For normalization, we spiked the gradient fractions with a protein marker prior to digestion and directly proceeded to MS analysis, i.e., without precipitation, in order to minimize the risk of losing low-abundance proteins.

Analysing bacteria in the early stationary phase of growth (OD_600_ of 2), i.e. our standard condition in several previous RBP studies (Holmqvist et al. 2016; Smirnov et al. 2016; Michaux et al. 2017; Holmqvist et al. 2018), we detected 2,555 *Salmonella* proteins in the original sample. Sedimentation profiles (gradient samples) were obtained for 2,225 proteins (Supplemental Table 1 and Fig. 1C), which is similar to the number of detected proteins in a recent RBP discovery study in *Salmonella* (Urdaneta et al. 2019). We observed a good correlation of overall protein abundance between the two gradients (Supplemental Fig. S2A-B), arguing that there would be few false positives resulting from potential RNase-induced protein insolubility. Permitting a ≤5-fold difference in protein intensity-based absolute quantitation (iBAQ, (Schwanhäusser et al. 2011)) between the treated and untreated gradient, we proceeded with 1,914 proteins (Supplemental Fig. S2B), which covered ~75% of the detectable *Salmonella* proteome and showed an expected enrichment of cytosolic localization (Fig. 1C). Importantly, the 98 proteins with clear RNase-induced shifts towards lighter fractions were strongly enriched in annotated RBPs including ribosomal proteins (Fig. 1C). Moreover, the MS data matched well the western blot profiles of individual proteins, e.g., CsrA, Hfq, ProQ, and GroEL (Fig. 1D), suggesting that the global proteomics-based GradR approach could be used to discover new RBPs.

Not all known RPBs shifted fractions upon RNase treatment (Fig 1C, Supplemental Fig. S2C-E). Obvious false negatives include the cold shock-like proteins CspA, CspC, CspE, and CspB (a.k.a. CspJ), which shifted only marginally (Supplemental Fig. S2E) despite the fact that each of them binds hundreds of different cellular transcripts (Michaux et al. 2017). These RBPs are difficult to assess; with the present glycerol concentration they are LMW to begin with (Supplemental Fig. S2E). Regarding false positives, proteins might shift in the gradient not because they themselves loose an RNA ligand but because they are in a complex with a *bona fide* RBP.

### Classification and visualization of RNase-induced changes

To classify RNase-dependent sedimentation profiles of proteins, we used two different descriptors: the number of fractions shifted, and the change in general presence and distribution in the gradient. We consider the latter as a rough indicator of the extent of a protein’s interactions with cellular transcripts which themselves vary greatly in length (thus, molecular weight) and shape. The result is a map with four quadrants (Fig. 2).

**Fig. 2.**
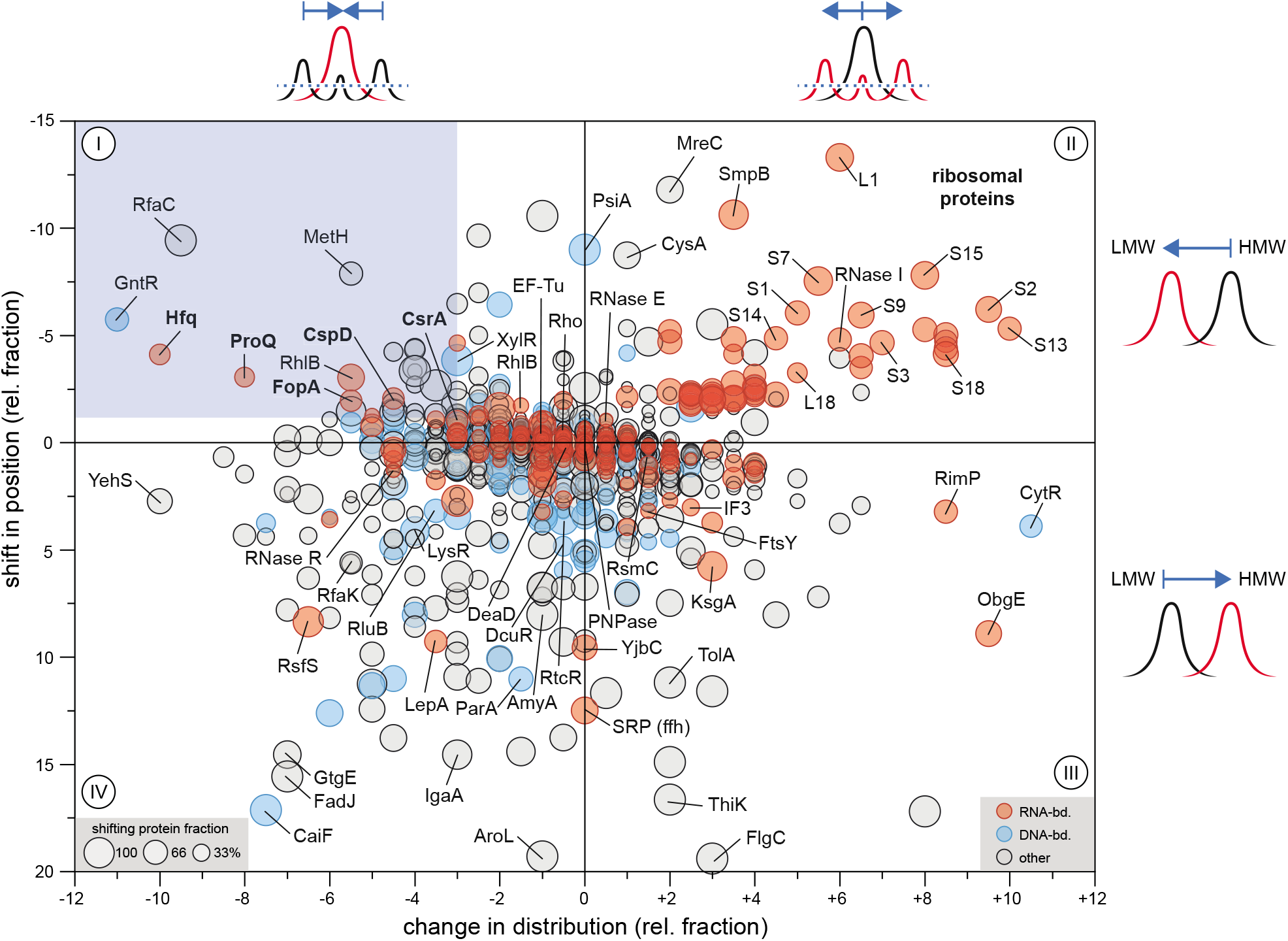
RBPs shift and reduce distribution in the gradient. In addition to a shift to LMW fractions, the presence and distribution of Hfq, ProQ, CspD, and CsrA was reduced in the gradient upon RNase treatment (quadrant I). Ribosomal proteins increased distribution in the gradient while still shifting to LMW fractions (II). Some proteins shifted to HMW fractions upon RNase treatment or pelleted (III, IV).

Quadrant I at the top left was enriched with well-established RBPs, e.g., CsrA, Hfq and ProQ. In general, the proteins in this quadrant each spread over several neighbouring fractions but their peak both sharpens and shifts towards lighter fractions upon removal of their RNA partners. We will describe further below how this quadrant unveiled a new FinO-domain RBP, but before this discuss several intriguing patterns in the other three quadrants, which represent opposing and more complex scenarios (Fig. 2).

### Sedimentation profiles of many Salmonella proteins are sensitive to RNase treatment

Quadrant II is dominated by r-proteins of the ribosome (Fig. 2), which is the largest known ribonucleoprotein particle (RNP) in *Salmonella* (Smirnov et al. 2016; Burley et al. 2018). 70S ribosomes typically sediment in high molecular weight (HMW) fractions 19-20 as well as the pellet (Supplemental Figs. S1A-B, S2C), but as RNase fragments ribosomal RNA, the 70S RNP decomposes and sub-complexes appear (Supplemental Fig. S1B). Concomitantly, r-proteins of the small (30S) and large (50S) subunits appear in LMW fractions and tend to spread over more fractions, i.e., increase in distribution (Fig. 3A).

**Fig. 3.**
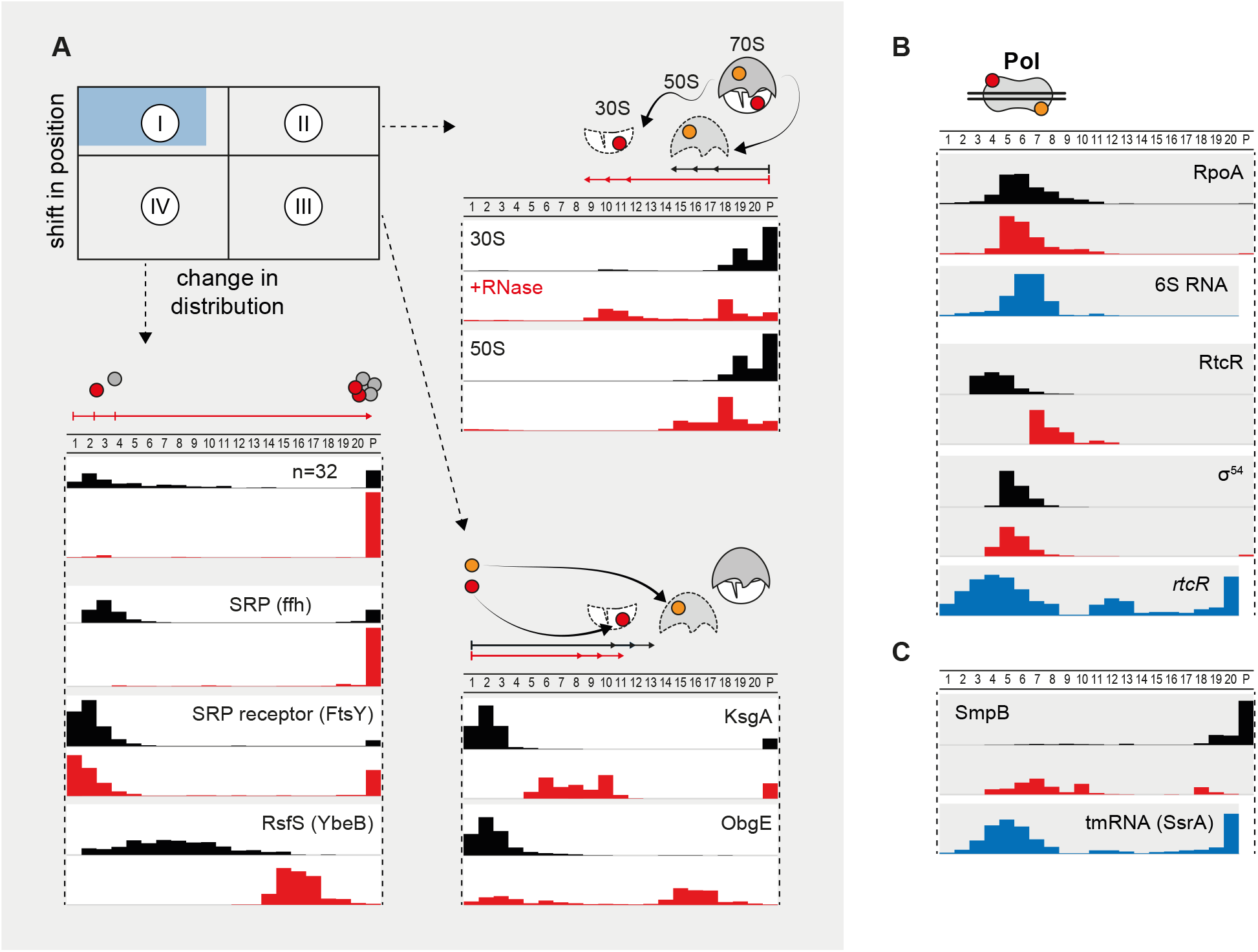
Complex scenarios are possible for ribosome or polymerase associated proteins. (A) Map from Fig. 2 indicating locations of protein candidates in further panels. In quadrant II, 70S ribosomes disintegrated and proteins shifted towards the LMW fractions corresponding to 30S and 50S fractions, increasing the distribution in the gradient. In quadrant III, translation factors occupied subunit fractions, shifting from LMW fractions to ribosomal fractions. In quadrant IV, a number of factors pelleted upon RNase treatment that may require RNA for integrity. An interesting example was the SRP protein, but also some transcription and translation factors were shifted to HMW fractions. (B) The RNAP complex did not shift. RNAP proteins correlated with 6S RNA (RNA sedimentation profiles are represented as blue bars) (Smirnov et al. 2016). The sigma S factor shifted to LMW fractions. Interestingly, transcription factors shifted between polymerase associated fractions, for example RtcR that was allocated before RNase treatment closely to *rtcR* RNA (Smirnov et al. 2016). (C) The trans-translation factor SmpB shifted to polymerase associated fractions after RNase treatment, and partly overlapped with its associated tmRNA (Smirnov et al. 2016).

In quadrant III, we note several proteins that normally occur in LMW fractions (in line with their molecular weight), but after RNase treatment show shift to larger complexes in HMW fractions and concomitantly broader distribution (Figs. 2, 3A). These include several ribosome-associated factors such as the 16S rRNA methyltransferase KsgA and the 50S GTPase ObgE, which upon RNase-treatment occupied the 30S or 50S fractions, respectively. KsgA and ObgE have been observed in cryo-EM reconstruction to interact with 30S and 50S ribosomes, respectively (Boehringer et al. 2012; Feng et al. 2014), and both were predicted to interact with rRNA in these structures. Many other proteins show similar behaviour (Supplemental Fig. S3), and these may be candidates for presently unrecognized functions in protein synthesis.

Quadrant IV represents proteins that shift to HMW fractions and become less spread upon RNase treatment; most of them end up in the pellet, which could mean either association with very large complexes (e.g. 70S) or simply, insolubility (Fig. 3A). The signal recognition particle (SRP), composed of protein Ffh and 4.5S RNA, stalls translating ribosomes until their recruitment by receptor FtsY to the membrane for co-translational translocation of the protein to be synthesized (Walter et al. 1981; Halic et al. 2004). The Ffh protein shifted almost completely to the pellet as its 4.5S RNA partner was digested. By contrast, several proteins did not pellet but appeared to associate with RNA polymerase (RNAP), as discussed in the next section, or appeared in other HMW fractions. For example, ribosome silencing factor RsfS shifted to 50S subunit fractions 15-17, which would be in line with its described molecular function as an antiassociation factor (Li et al. 2015) that targets the large ribosomal subunit during starvation (Fig. 3A).

### RNase induces shifts of proteins into and out of RNAP fractions

The RNA polymerase (RNAP) is a ~450-kDa complex comprised of the α/β/β’/ω (RpoA/B/C/Z) subunits, and σ factors that control promoter binding. In addition, RNAP is an RNP when in complex with 6S RNA (Wassarman and Storz 2000). While RNAP itself did not shift upon RNase treatment, several transcription factors (TFs) changed distribution (Fig. 3B). One striking example is the TF RtcR, which together with σ^54^ controls transcription of the RNA repair system RtcAB (Genschik et al. 1998). RtcR sedimented in the same fractions (4-6) as did its own mRNA (Smirnov et al. 2016); these fractions partly overlap with RNAP. Yet, RNase treatment shifted RtcR to HMW fractions 7-9 (Fig. 3B). It is tempting to speculate that binding to its own mRNA modulates the interaction of RtcR with RNAP, and more generally, that RNA-dependent associations with RNAP could provide specific feedback mechanisms in transcriptional regulons (see discussion in (Holmqvist and Vogel 2018)).

Another prominent RNase-induced shift towards RNAP fractions is seen with SmpB, the protein in the tmRNA-containing RNP that rescues stalled ribosomes (Rae et al. 2019). Interestingly, tmRNA was previously observed to partially sediment in the RNAP fractions as well (Smirnov et al. 2016), indicating a potential link between translation and transcription of tmRNA (Fig. 3C).

### A candidate FinO-domain RBP

Quadrant I (Fig. 2) represents profiles most expected for a typical RBP, i.e., RNase treatment would induce a shift towards LMW fractions and cause a more compact distribution in the gradient. After hierarchical clustering of sedimentation profile changes, we obtained two main clusters of proteins with RNase-driven enrichment in either fraction 1 or fraction 3 (red, blue, respectively, constrains: shift <-1, change in distribution <-3, shifting protein fraction >40%, Fig. 4A). The former cluster contained not only CspD, Hfq, and ProQ, but also the FopA protein of unknown function. As per the global MS data, FopA primarily sedimented in fractions 2-3, but shifted to fractions 1-2 upon RNase treatment (Fig. 4B). Western blot analysis using a *Salmonella* strain in which we had fused a C-terminal 3×FLAG tag to the *fopA* gene confirmed this prediction (Fig. 4C), and made FopA an attractive candidate for a previously unrecognized RBP.

**Fig. 4.**
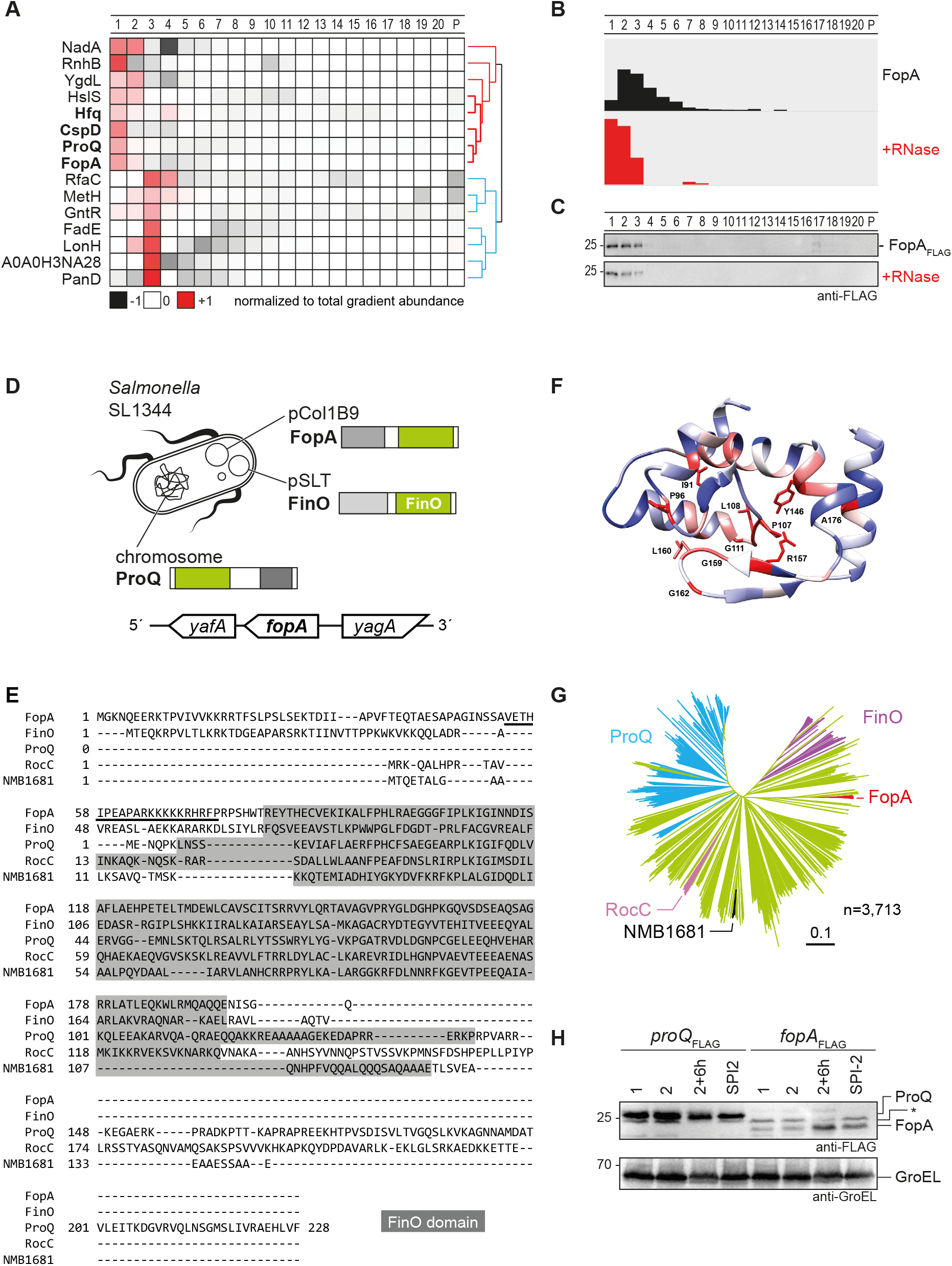
The FinO-domain protein FopA is abundant with a unique N-terminal domain. (A) Hierarchical clustering of candidates that shifted to LMW and reduced distribution (Fig. 2, blue box, filtered to >40% shifting protein fraction) resulted in clustering of Hfq, ProQ, CspD, and the RBP candidate FopA in the red cluster. In the blue cluster proteins shifted towards fraction 3. (B) The MS quantification of FopA showed a similar sedimentation shift as for ProQ, which was confirmed by Western blot analysis (C). (D) *Salmonella* harbours three FinO-domain proteins. ProQ is encoded on the chromosome whereas FinO and FopA are encoded on plasmids. (E) FopA homologs show a unique N-terminal domain that is different to the FinO N-terminal domain. The central FinO domain in FopA (PF04352, grey background) shows conserved key residues. (F) Homology model of the FinO domain of FopA with color-coded conservation (red indicates high conservation). (G) FopA proteins represent a distinct branch (red) in a phylogenetic analysis based on protein sequences of 2,569 FinO-domain containing proteins (PF04352, Pfam 32.0, Clustal Omega analysis). (H) FopA levels are strongly elevated at late stationary phase and increased in SPI-2-inducing conditions. * indicates anti-body cross-reaction.

FopA is encoded on the resistance plasmid pCol1B9 (a.k.a. P2 of incompatibility group IncIα, GenBank: HE654725.1), which is one of three plasmids in the *S. enterica* strain SL1344 used here (Fig. 4D) (Asano et al. 1999). pCol1B9 expresses colicin Ib, a narrow-spectrum bacteriocin against *Enterobacteriaceae*. The *fopA* gene lies between *yafA* and *yagA*; all of these genes encode proteins of unknown function. Intriguingly, according to a recent update of the Pfam database (El-Gebali et al. 2018), FopA carries a FinO domain (entry: PF04352, Fig. 4E), making it the third such protein in *Salmonella*, in addition to FinO from the conjugative transfer locus on plasmid pSLT, and ProQ from the chromosome. Moreover, FopA possesses the conserved FinO domain residues Arg157 and Tyr146, which are essential for RNA binding by ProQ (Pandey et al. 2020), and key residues in a homology model of FopA (Fig. 4F). Apart from that, FopA represents a distinct branch of FinO-domain proteins (Fig. 4G), with plasmid or chromosome encoded homologs in *Escherichia, Shigella*, and *Klebsiella*. Specifically, FopA lacks the N-terminal domain of FinO and the C-terminal domain of ProQ but exhibits a distinct N-terminal domain with two stretches of positively charged residues.

Available RNA-seq data (Canals et al. 2019) predict differential expression of *fopA* under specific conditions. In western blot analysis FopA accumulated in stationary phase (2+6h), but also in media that induce the *Salmonella* pathogenicity island 2 (SPI-2) (Fig. 4H). By semi-quantitative comparison with ProQ for which a copy number of 600-1,200 monomers per *E. coli* cell has been determined (Wiśniewski and Rakus 2014; Soufi et al. 2015) (Supplemental Fig. S4A), we infer that FopA is present in 200-400 copies per *Salmonella* cell (stationary phase). In other words, FopA is an abundant candidate RBP with an intracellular concentration in the micro-molar range.

### FopA is a plasmid-encoded RBP with a unique target suite

To test RBP activity of FopA and identify potential RNA ligands *in vivo*, we performed a RIP-seq analysis (Chao et al. 2012), sequencing transcripts after coimmunoprecipitation (coIP) with the tagged protein in the *fopA*::3×FLAG strain grown to early stationary phase. Comparison with previous RIP-seq results for ProQ obtained in the same growth phase (Smirnov et al. 2016) yielded two key observations. Firstly, despite their sharing a FinO domain, FopA and ProQ have very different target suites (Fig. 5A). Reads obtained with FopA are dominated by Inc (Fig. 5A-C), which is a ~70-nt regulatory RNA expressed from the same plasmid that encodes FopA. By contrast, Inc is hardly recovered in RIP-seq of ProQ (Smirnov et al. 2016) (reanalysed in Fig. 5A).

**Fig. 5.**
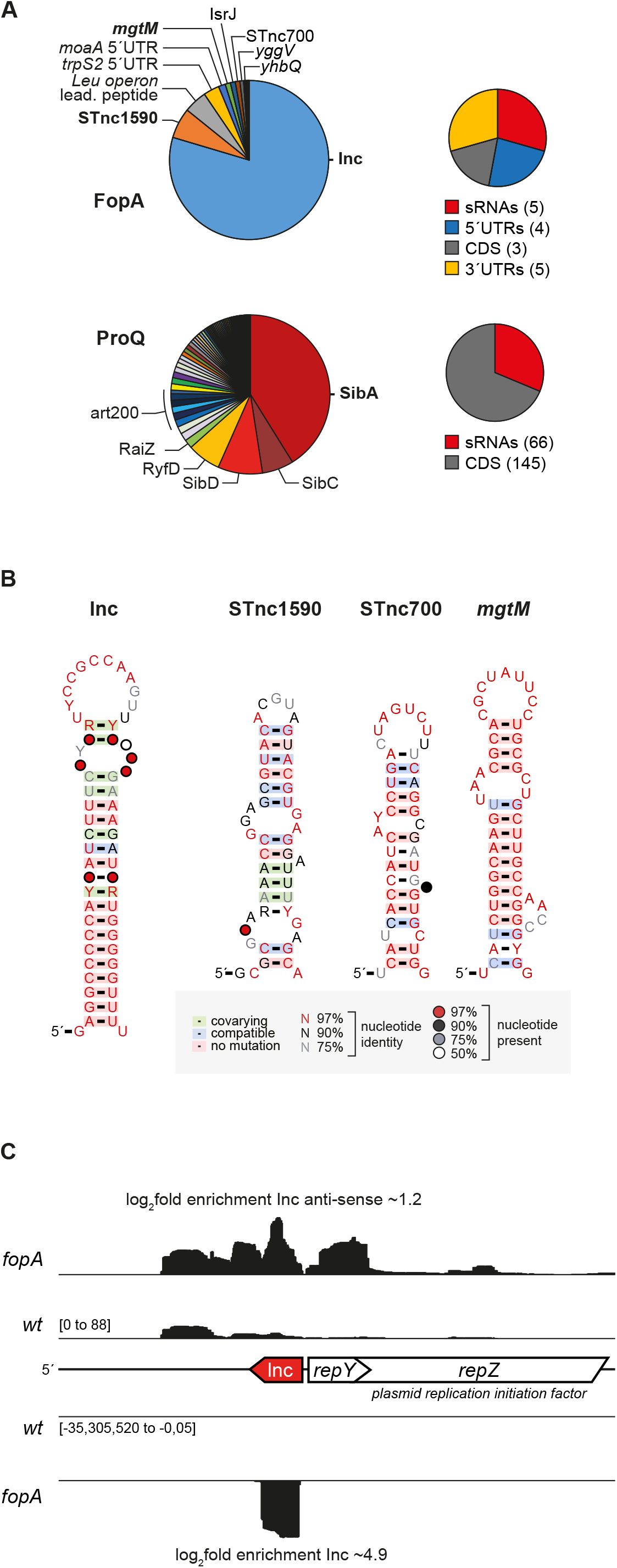
FopA is a global RBP that targets structured RNAs. (A) The major targeted RNA was the antisense RNA lnc, followed by STnc1590, *mgtM*, and STnc700 (log2-fold change > 4, n = 17). The RNA targetome of FopA was very different to ProQ recovered RNAs that were reanalysed from (Smirnov et al. 2016). (B) FopA targeted RNAs were analysed for conservation motifs by the CMfinder algorithm. Homologs were searched by GLASSgo (1.5.0, Freiburg RNA tools, (Lott et al. 2018)). The CMfinder algorithm determined RNA motifs (0.4.1.18, (Yao et al. 2005)), and R2R was used for visualization (1.0.6, (Weinberg and Breaker 2011)). (C) FopA-coIP read-coverage of the *repZ* locus. Inc is heavily enriched, together with the sparsely expressed *repZ* mRNA.

Secondly, although their targets are distinct in primary sequence, FopA and ProQ both associate with highly structured transcripts (Fig. 5B). Interestingly, Inc and STnc700 both end with a 4U stretch, i.e., which is a recently proposed 3’ end-located recognition element for binding by the FinO-domain (Stein et al. 2020). As previously observed with ProQ (Holmqvist et al. 2018), the targets of FopA share no obvious primary sequence or structural motif as none could be predicated that fits all the main targets (Supplemental Fig. S4B), hence we conclude that FopA binds an unknown hidden structured RNA code.

Inc is a *cis*-antisense RNA in the 5’ region of *repZ* encoding the replication initiator protein of pCol1B9 (Asano et al. 1998). The low-abundance *repZ* mRNA is also enriched by coIP with FopA (Fig. 5C). Interestingly, Inc/*repZ* and FopA show equal phylogenetic distribution, indicating functional linkage (Supplemental Fig. S4C). While Inc and *repZ* are plasmid-encoded transcripts, other top targets of FopA are transcribed from the *Salmonella* chromosome. These latter targets include STnc1590, which is a *cis*-antisense RNA to STnc1580 (found in several different enterobacteria); the non-coding RNA STnc700 from the leader region of the histidine operon mRNA; and the transcribed *mgtM* region that is important for ATP-sensing and regulation of the *mgtCBR* operon (Lee and Groisman 2012). Interestingly, the *mgtCBR* operon is also regulated by ProQ (Westermann et al. 2019). In summary, FopA is the third FinO-domain protein of *Salmonella*, with targets in both a plasmid and the core genome of this bacterium.

### FopA accelerates RNA duplex formation

Seeking to confirm the predicted RBP activity of FopA, we selected the Inc-*repZ* RNA pair for *in vitro* binding experiments. These two RNAs form a kissing complex that subsequently progresses to a four-way junction (reviewed in (Kolb et al. 2001)). For our binding experiments, we purified recombinant *Salmonella* FopA protein after expression in *E. coli* and rendered it RNA-free by ion-exchange chromatography, thus reaching a purity of more than 95 % (Fig. 6A, Supplemental Fig. S5).

**Fig. 6.**
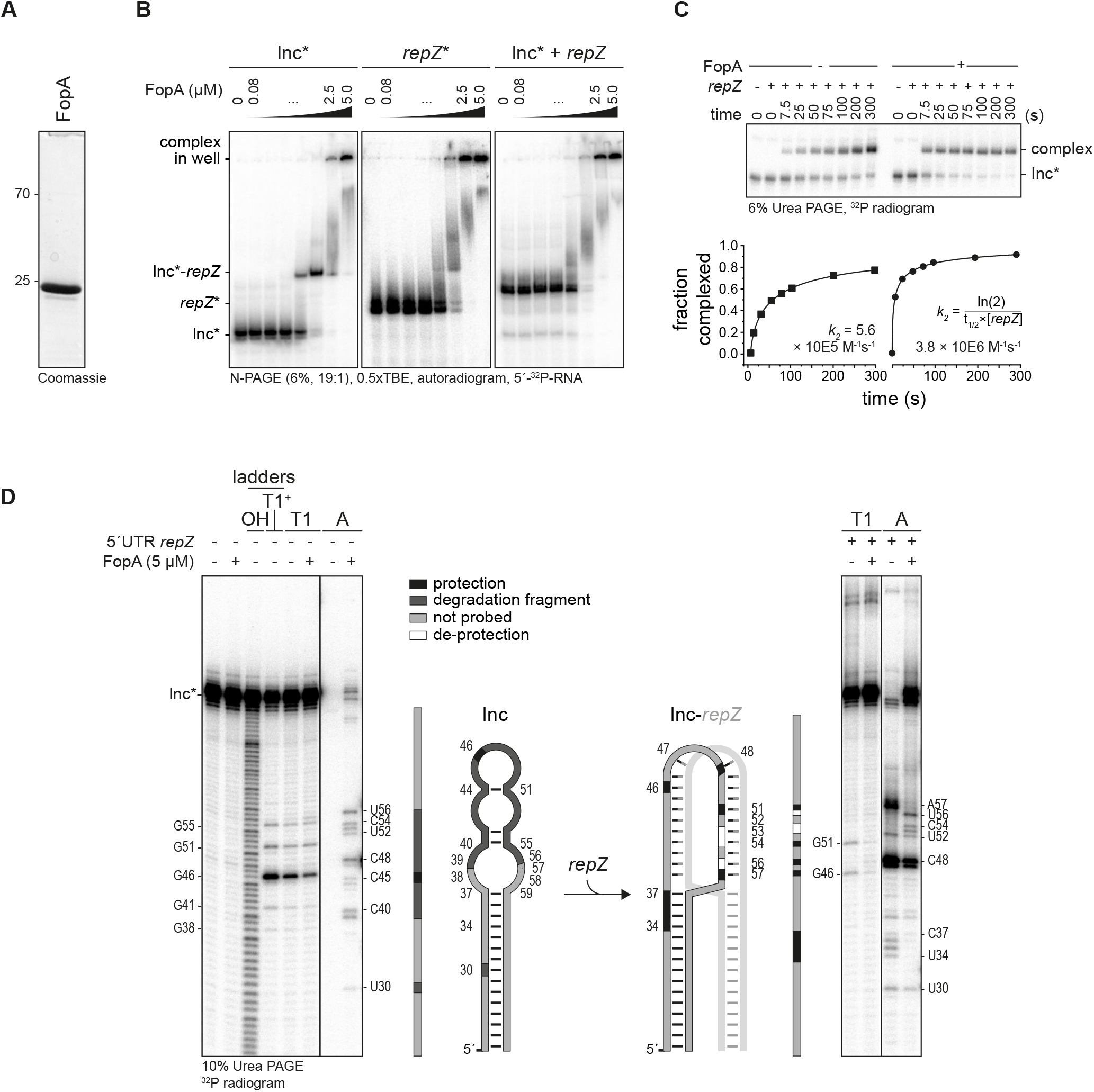
FopA protects Inc and anneals it to the 5’UTR of *repZ*. (A) Purified FopA with cleaved-off tag. (B) FopA bound Inc, *repZ*, and the Inc-*repZ* complex with an apparent *K_D_* about 1 μM. n=2 (C) FopA accelerated Inc*-*repZ* complex formation. *k*_2_ represents the rate constant. n=2 (D) FopA protected full-length Inc alone and in the complex with *repZ* from RNase A degradation at loop regions. n=2

Using electrophoretic mobility shift assays (EMSA), we determined an apparent affinity constant of ~1 μM for FopA and Inc RNA (Fig. 6B), which is a good fit with both the intracellular concentration of FopA and reported affinities of other FinO-domain proteins (Attaiech et al. 2016; Smirnov et al. 2016; Holmqvist et al. 2018; Bauriedl et al. 2020). FopA displayed similar affinities for the *repZ* 5’UTR fragment fully antisense to Inc or for a preformed Inc-*repZ* RNA complex (Fig. 6B). Since FopA bound Inc and *repZ* both individually and as a complex, we also assessed whether it affects the kinetics of Inc-*repZ* complex formation. Once formed, the Inc-*repZ* complex has no apparent off-rate, hence the initial association rate is equivalent to its formation rate (*k*_2_, as previously established for Inc and orthogonal systems (Persson et al. 1988; Tomizawa 1990; Asano et al. 1998)). Importantly, FopA accelerated complex formation of the Inc and *repZ* RNAs by ~7-fold (Fig. 6C). As such, the primary function of FopA may be that of an RNA chaperone, similar to the molecular function of its plasmid-encoded sibling FinO in the FinP-*traJ* conjugation control system (Arthur et al. 2003).

To map FopA binding sites within Inc, we performed RNA structure probing with the single-strand specific endoribonucleases, RNase A (cuts between unpaired pyrimidines) and RNase T1 (cuts at unpaired guanosines). As shown in Fig. 6D, FopA protected Inc from cleavage in the loop region at G_46_, and in general, protected Inc from complete degradation by RNase A. In the Inc-*repZ* complex, the inter-bridged region was protected between U_34_-U_37_ and at A_57_. Concurrently, Inc was deprotected at positions U_53_, C_54_, and U_56_, indicating an effect on the RNA structure at the cross-junction point. Taken together, by showing binding of FopA to its major RNA ligands *in vivo*, we established proof-of-principle that GradR can predict a previously unrecognized bacterial RBP.

## DISCUSSION

GradR is conceptually related to two recently published approaches for eukaryotes, R-DeeP and DIF-FRAC, which used sucrose instead of glycerol gradients (Caudron-Herger et al. 2019) or size-exclusion chromatography (Mallam et al. 2019). Together, these two latter studies predicted >1,400 so-called RNA-dependent proteins or complexes in human cells; in other words, >7% of the human proteome would be sensitive to RNase treatment, which is in the same range as the >5.5% predicted here for a bacterial proteome. In order to visualize these RNase sensitivities and evaluate individual proteins in the context of their physiological role and published literature, we provide an interactive browser (Fig. 7, www.helmholtz-hiri.de/en/datasets/GradRSeT user: reviewer, pw: welcome) that also allows one to quickly compare proteins from related organisms via Pfam protein domain annotations (El-Gebali et al. 2018).

**Fig. 7.**
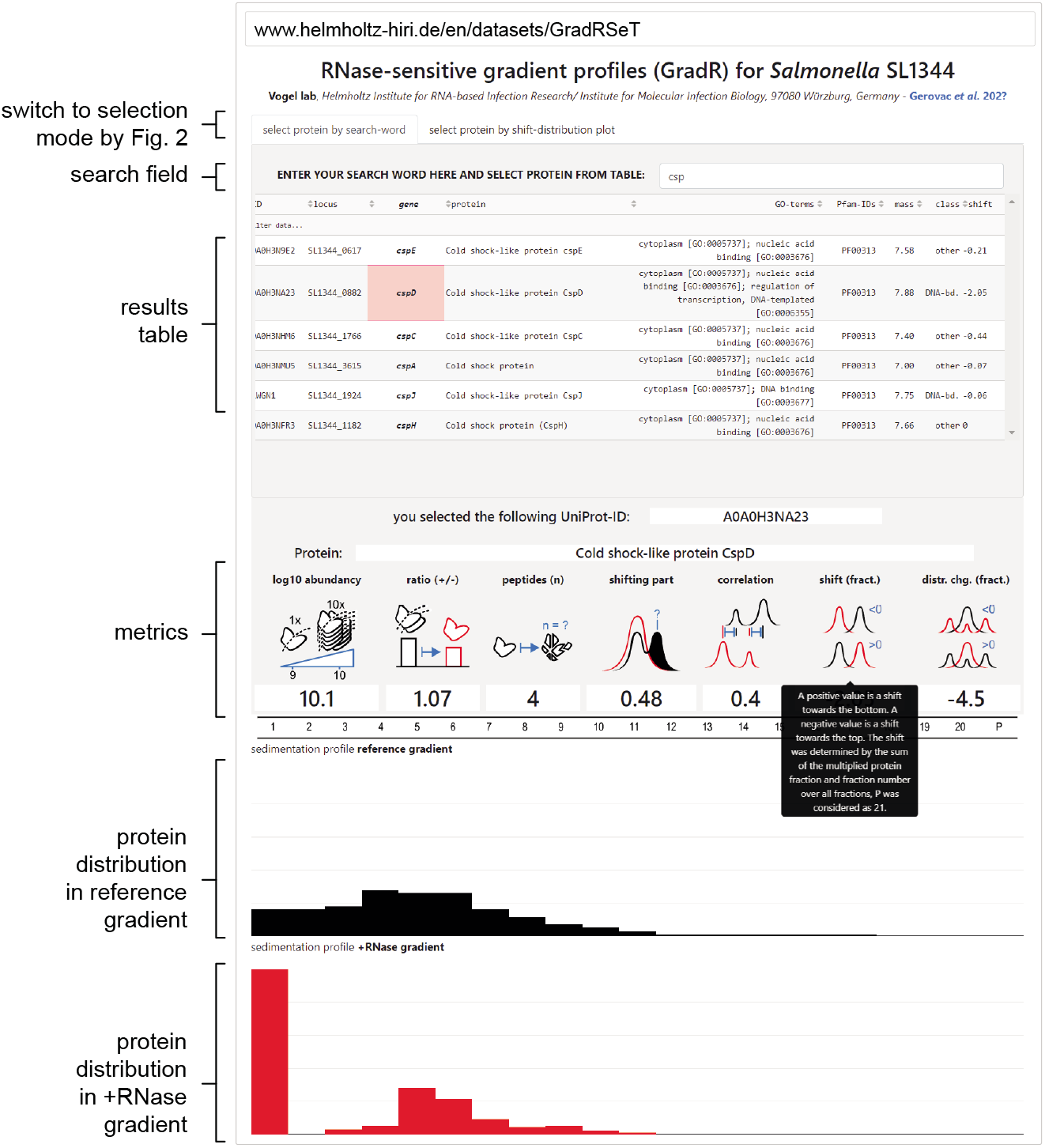
Visualizing sedimentation patterns and RNase sensitivity data. Complex GradR data can be visualized and explored in a browser, available at www.helmholtz-hiri.de/en/datasets/GradRSeT.

There has been a flurry of new approaches using organic extraction or silica-based purification to globally predict RBPs, several of which also established proof-of-principle for usability beyond eukaryotes with *E. coli* or *Salmonella* (Queiroz et al. 2019; Shchepachev et al. 2019; Urdaneta et al. 2019). We cross-compared our *Salmonella* GradR data set with the list of *Salmonella* RBP candidates predicted by Phenol Toluol extraction (PTex, (Urdaneta et al. 2019)). PTex predicted 172 RBP candidates, of which 29 had an annotated GO-term ‘RNA-binding’ representing ~21% of all *Salmonella* proteins within that GO term. For comparison, GradR detects 210 RNase-sensitive proteins, 34 of which (~25%) have the GO term ‘RNA binding’. Interestingly, the GradR and PTex predictions share only 24 RBP candidates, 62% of which are established RBPs (Supplemental Fig. S6A). As far as cross-comparison of *Salmonella* with *E. coli* data permits, we correlated GradR with silica-based recovery of UV-crosslinked RBP candidates to RNA by TRAPP (Shchepachev et al. 2019). Of the 377 candidates from TRAPP, GradR found 42; again, most (67%) of these shared proteins are established RBPs (Supplemental Fig. S6B). It is possible that the crosslinking step used in these other methods (Asencio et al. 2018; Queiroz et al. 2019; Shchepachev et al. 2019; Trendel et al. 2019; Urdaneta et al. 2019) helps to enrich weakly interacting candidates. By contrast, GradR requires RNPs to remain stable after lysate preparation. Therefore, we consider GradR as an orthogonal RBP discovery method to UV-crosslinking based approaches. Most importantly, its generic nature provides an experimental route to glance at the major RBPs of any microbe of interest, which becomes important in light of the thousands of understudied bacteria now known to populate the human body.

*Salmonella* being a well-characterized pathogen, potential RBPs amongst its virulence factors are of special interest. While previous RNA-binding domain based searches failed to predict RBPs amongst *Salmonella’s* secreted virulence factors (Sharan et al. 2017; Tawk et al. 2017), we note that the virulence-associated protein YopD of the related pathogen *Yersinia pestis* is well-known to moonlight as an RBP (Chen and Anderson 2011). In addition, *Listeria monocytogenes* has just been shown to secret an RBP to manipulate infected host cells (Gerovac and Vogel 2019; Pagliuso et al. 2019). Interrogating RNase sensitivity of *Salmonella* virulence-associated proteins (Supplemental Fig. S7A), we observe pronounced shifts towards central fractions for the invasion protein InvF, the secretion system apparatus protein PrgJ, and the protein encoded by *orf319*. The invasion protein OrgB, the bacteriophage-encoded virulence factor GtgE, effector protein SseK1, and an uncharacterized prophage-encoded NTPase (SL1344 2214) pelleted upon RNase treatment. Hypothetical virulence protein MsgA, sedimenting in fractions 1-3, shifted to fraction 1 upon RNase treatment, making it a good RBP candidate. Importantly, GradR might find even more potential RBPs when *Salmonella* is grown under conditions that fully induce its virulence genes, so more proteins accumulate to levels that permit detection by MS.

Other classes of proteins lend themselves for inspection in GradR data. For example, studies in eukaryotes have accumulated evidence that many enzymes from intermediary metabolism may moonlight as RBPs, often by binding their own mRNAs (Hentze et al. 2018). Of studies in bacteria, one in *E. coli* proposed ~1,000 RBP candidates that comprised mainly metabolic proteins (Shchepachev et al. 2019). A primary example of a metabolic RBP in *E. coli* is the iron-sensing protein Aconitase B. This enzyme acts in the tricarboxylic acid (TCA) cycle and directly stabilizes its own mRNA (*acnB*) when intracellular iron becomes scarce (Benjamin and Massé 2014). Here, the 93.5 kDa AcnB protein is present in fraction 3 in a single peak that is unaffected by RNase treatment (Supplemental Fig. S7B,C). Potential reasons for failure to detect this established RBP by GradR include the use of iron-rich media in our study, and a possible instability of the AcnB-mRNA complex in lysate. However, we do observe shifts upon RNase treatment such that several metabolic proteins move to HMW fractions (AraA, AmyA, CbiK, RfaG) or the pellet (RfaK, NarG, FrdB, CyoB, FadJ, AroL), which may indicate that certain metabolic enzymes require intracellular association with RNA for integrity.

The power of GradR is well illustrated by its ability to validate not only all three sRNA-related RBPs (CsrA, Hfq and ProQ) of *Salmonella* but also to predict a new FinO-domain RBP from a plasmid, i.e., a common type of extrachromosomal element for which there has been almost no systematic annotation effort (Pilla and Tang 2018). FinO-domain proteins are a growing class of RBPs that target structured RNA molecules in the cell, distinct from tRNA and rRNA (Attaiech et al. 2016; Smirnov et al. 2016; Holmqvist et al. 2018; Bauriedl et al. 2020; Melamed et al. 2020). While most of the recent attention to this RBP class has been on global post-transcriptional regulation by the chromosomally encoded ProQ and RocC proteins, the class’ founding member—FinO— just like FopA is encoded on a plasmid (Timmis et al., 1978). FinO’s assumed main cellular RNA targets (the FinP sRNA-*traJ* mRNA pair) are also plasmid-encoded (Mark Glover et al. 2015), as is FopA’s main target, the Inc RNA. By contrast, ProQ stably associates with transcripts from the chromosome. Nonetheless, all these RBPs select their very different targets from the same pool of cytoplasmic transcripts, with the same type of FinO domain. Importantly, since the *Salmonella* virulence plasmid pSLT encodes a homolog of *E. coli* FinO (Fig. 4d), our work establishes this bacterium to express all three such RBPs (FinO, ProQ, FopA). *Salmonella* could thus be a test bed for domain swapping experiments to address what intramolecular or cellular factors, including intracellular localization, determine target selection. In addition, the RNA interaction by ligation (RIL)-seq technique (Melamed et al. 2016), which in *E. coli* revealed diverse overlapping, complementary, or competing roles for ProQ and Hfq (Melamed et al. 2020), may offer a more sensitive tool than coIP to study currently unknown RNA interactions amongst the target suites of these three FinO domain RBPs.

Structural and molecular progress on defining its RNA interface notwithstanding (Ghetu et al. 2000; Ghetu et al. 2002; Arthur et al. 2011; Attaiech et al. 2016; Immer et al. 2018; Pandey et al. 2020), the structure of a FinO domain with a bound RNA remains to be solved. The 22 kDa FopA protein can be purified to homogeneity (Fig. 6a) and its RNA ligands offer tight folds, both of which is conducive to structural studies. Given FopA’s distinct sequence (Fig. 4E,G), we expect a structurally resolved FopA–RNA complex to provide important complementary information in the quest for the rules that govern the intriguing selectivity of the FinO domain for RNA structure.

## ACKNOWLEDGEMENTS

We are grateful to Barbara Plaschke for technical assistance; Alexander Westermann, Jens Hör, Andreas Schlosser, Stephanie Lamer, and all lab members for helpful discussions; Benedikt Beckmann, Cynthia Sharma and Gisela Storz for providing comments on the manuscript.

## AUTHOR CONTRIBUTIONS

M.G. and J.V. designed experiments. M.G. performed all experiments unless stated otherwise. M.G. and J.K. purified protein, C.K. provided resources. M.G. performed and L.B. validated data analysis. M.G. programmed the GradR browser. M.G. and J.V. wrote the paper with input from Y.E.M. All authors approved the final version.

## MATERIALS & CORRESPONDENCE

should be addressed to J.V.

## METHODS

### Cell lysis, RNase digestion, and gradient fractionation

*Salmonella* SL1344 cells were grown in Lysogeny broth (LB) media to transition phase at OD_600_ of 2.0, pelleted at full speed, and frozen at −20 °C. In total 60 OD of cells were lysed in 500 μl lysis buffer (20 mM Tris/HCl pH 7.5, 150 mM KCl,10 mM MgCl_2_, 1 mM DTT, 2 mM PMSF) by glass beads (0.1 mm, BioSpec Products) in the Retsch MM200 at 30 Hz for 10 min at 4 °C. The cell lysate was cleared at full speed for 10 min and the supernatant was transferred. 20 μl RNase A/T1 mix (2 μg μl^−1^ / 5 U μl^−1^, Thermo Scientific) was added to 100 μl lysate (OD_260_ ~ 150) for RNase digestion for 20 min at 20 °C. The reaction was stopped on ice, diluted with 1 volume lysis buffer, and loaded completely onto a 10-40% glycerol (w/v) gradient in lysis buffer. The proteins were sedimented in the SW40Ti rotor (Beckman Coulter) at 100,000 × g for 17 h at 4 °C. The gradient was fractionated in 20 equal samples (~600 μl) from the top and the pellet was resuspended in an additional fraction. RNA was extracted by PCI for RNA (X985.3, Carl Roth) for RNA integrity control gels. Protein samples were prepared for SDS-PAGE analysis and western blotting.

We analysed gradients with and without 0.2% Triton X100 in the lysis and gradient buffers (Supplemental Fig. S2G-I). Detergents solubilize membrane-associated proteins and prevent unspecific interactions, but also interfere with peptide recovery and mass spectrometric analysis. Both GradR experimental condition replicates were well correlated (Supplemental Fig. S2G-I, for details see section about data analysis) and we present here data of the GradR experiment without Triton X100 and complete insolution MS sample preparation.

### In-solution MS sample preparation

0.2 μg proteomics dynamic range standard set (UPS2, Sigma-Aldrich) was added as spike-in to 50 μl protein samples of each fraction. Samples were diluted with two volumes denaturation buffer (100 mM Tris/HCl pH 8.5, 12 M urea). 5 mM tris(2-carboxyethyl)phosphine (TCEP, Bond-Breaker, neutral pH, Thermo Scientific) was added and samples were incubated at room temperature for 20 min. 10 mM iodoacetamide (IAA) was added and samples were incubated at room temperature for 15 min in the dark. 0.25 μg Lys-C protease (Promega) was added and incubated for 4 h at 37 °C. Samples were diluted with three volumes digestion buffer (100 mM Tris/HCl pH 8.5, 1 mM CaCl_2_) and 0.25 μg Trypsin (Sequencing Grade, Promega) were added and incubated o/n at 37 °C. 5% formic acid (FA) was added for acidification and samples were cleared by centrifugation at full-speed for 10 min. Acidified sample supernatant was loaded onto methanol activated stage-tips (C18) by centrifugation for 5-10 min at 2,000 × g. Tips were washed three times with washing solution (2% acetonitrile (ACN), 0.3% trifluoroacetic acid (TFA)) and eluted in 0.5 ml protein-low bind tubes with two times 20 μl elution solution (60% ACN, 0.3% FA). Samples were snap-frozen in liquid nitrogen and lyophilized. For liquid chromatography analysis, samples were solubilized in application solution (2% ACN, 0.1% FA), sonicated, and 12 μl transferred to HPLC tubes.

### NanoLC-MS/MS Analysis and MS analysis

Samples were MS analysed in the lab of Andreas Schlosser by Stephanie Lamer. NanoLC-MS/MS analyses were performed by an Orbitrap Fusion (Thermo Scientific) equipped with a PicoView Ion Source (New Objective) and coupled to an EASY-nLC 1000 (Thermo Scientific). Peptides were loaded on capillary columns (PicoFrit, 30 cm × 150 μm ID, New Objective) self-packed with ReproSil-Pur 120 C18-AQ, 1.9 μm (r119.aq., Dr. Maisch). Samples were analysed by a 120 min linear gradient from 3-40% acetonitrile and 0.1% formic acid at a flow rate of 500 nl/min. MS and MS/MS scans were acquired in the Orbitrap analyser with a resolution of 60,000 and 15,000, respectively. Higher-energy collisional dissociation (HCD) fragmentation was applied with 35% normalized collision energy. We used top speed data-dependent MS/MS method with a fixed cycle time of 3 s. Dynamic exclusion was applied with a repeat count of 1 and an exclusion duration of 60 seconds (singly charged precursors were excluded from selection). Minimum signal threshold for precursor selection was set to 50,000. Predictive AGC was used with AGC a target value of 2E05 for MS scans and 5E04 for MS/MS scans. EASY-IC was used for internal calibration.

Raw MS data files were analysed with MaxQuant version 1.6.2.2 (Cox and Mann 2008). Database search was performed with Andromeda, which is integrated in the utilized version of MaxQuant. The search was performed against the UniProt *Salmonella* Typhimurium UP000008962 (strain SL1344) and a database containing the proteins of the UPS2 proteomic standard. Additionally, a database containing common contaminants was used. The search was performed with tryptic cleavage specificity with three allowed missed cleavages. Protein identification was under control of the false-discovery rate (FDR, 1% on protein and peptide level). In addition to MaxQuant default settings, the search was additionally performed for the following variable modifications: Protein N-terminal acetylation, glutamine to pyro-glutamic acid formation (N-term. glutamine) and oxidation (methionine). Carbamidomethyl (cysteine) was set as fixed modification. For protein quantitation, the iBAQ intensities were used (Schwanhäusser et al. 2011). Proteins that could not be distinguished by peptides were listed individually.

### Data analysis

Relative protein abundance in gradient fractions were calculated by iBAQ. Absolute abundances were estimated by correcting for differences in digestion and C18 purification efficiency in each gradient fraction, through normalization to human albumin (Supplemental Table S1, norm. to spike-in, P02768ups|ALBU_HUMAN_UPS Serum albumin, chain 26-609, Supplemental Fig. S2A) that was added in equal amounts to each fraction as part of the UPS2 standard (Sigma-Aldrich). We selected albumin for normalization as the highest number of peptides (74) were recovered for it of all spike-in proteins. The protein ratio between reference and +RNase gradients was determined by the normalized total absolute protein abundance per gradient. For bar-diagram representation, the abundances for each individual protein were transformed into distributions across the gradient by dividing the abundance in each gradient fraction by the total abundance across the gradient (Supplemental Table S1, norm. to grad.). For visualization purposes, the y-axis of individual protein gradient profile plots were scaled to the largest value in either the reference or RNase treated gradient.

Relative protein level changes (d) per fraction (Fig. 4A) were determined by subtraction of the reference protein abundances from the RNase treated normalized protein abundances in each gradient fraction. These relative protein level changes per fraction were used for hierarchical clustering. The fraction of shifted protein was the sum of all positive relative protein level changes.

The mean *position* of a protein in a gradient was calculated as the mean of proportions of protein times each fraction number (pellet equals 21). The *shift* in protein distribution was calculated to capture the difference in relative protein positions between the reference and the RNase treated gradients (Supplemental Fig. S2A). Protein proportions were converted to binary vectors at two thresholds: 0.5% (to capture low abundance presence), and 2.5% (to capture more robust presence in a fraction). The Hamming distance was calculated for each protein between the reference and RNase-treated gradient at each threshold, providing a measure of similarity in protein *presence and distribution* profiles. The *change in presence and distribution* was then calculated as the mean of the Hamming distance at these two thresholds (Supplemental Fig. S2A).

Calculations were conducted in Microsoft Excel (Supplemental Table S1). All normalization operations are implemented in the Excel spread sheet and can be used for analysis of new datasets. Single-linkage hierarchical clustering on Euclidean distances was performed in Orange (3.23, University of Ljubljana, (Demsar et al. 2013)). Thresholds were applied as indicated in the panels through the *select row* widget. The *distance* widget calculated the distances between rows, and hence individual proteins, and the *hierarchical clustering* widget yielded clusters in continuous order that were visualized as a *heat map* in Orange or Excel with color-coding that represented the shifting protein fraction from −1 to +1 normalized relative protein levels per gradient.

Scatter plots were assembled in Origin (OriginLabs) as bubble charts. Classification of proteins was achieved through consecutively requesting terms from protein names and full GO term annotation (without case sensitivity, from UniProt, UP000008962, release 12/2019, (The UniProt Consortium 2018)) in Microsoft Excel (“RNA”, “riboso”, “transla”, and “nuclease” for the class *RNA-binding*; “DNA”, “transcript”, and “plasmid” for the class *DNA-binding*; “nucleo”, “metal”, “zinc”, “magnesium”, “iron”, “ATP, and “GTP” for the class *metal- and nucleotide-binding; metabolism* class proteins were derived from the Kyoto Encyclopaedia of Genes and Genomes (KEGG) database (sey00001, A09100 Metabolism, and subgroups); proteins annotated as both *metal- and nucleotide-binding* and *metabolic* proteins were extracted and are indicated as an isolated shared class; “membrane” for the *two times* class; “uncharacterized” for the *uncharacterized* class; proteins that remained unclassified were classified as *others*, or if only RNA- and DNA-binding proteins were shown, all others were classified as *others*). All other classifications are listed in Supplemental Table S4.

All figures were assembled in Adobe Illustrator. Pearson correlation coefficients between individual gradients or sedimentation profiles of proteins were calculated in Excel with the function ‘CORREL’ considering norm. to gradient protein abundance. We observed well correlated sedimentation profiles for the reference gradient with Grad-seq profiles (Supplemental Fig. S2F). In addition, the non-shifting proteins were highly correlated between reference and RNase treated gradients in experimental condition replicates with (.0) and without (.1) 0.2% Triton X100 (Supplemental Fig. S2G-I).

### Genomic 3×FLAG-tag labelling of *fopA* in *Salmonella enterica* Typhimurium SL1344

Constructs for recombination were amplified by PCR from genomic DNA with 35-45 nt 5’ and 3’ overhangs that were homologous to the recombination sites of the gene of interest. Wild-type *Salmonella* strain (JVO-1574) was transformed with pKD46 (Datsenko and Wanner 2000), selected by ampicillin resistance on a plate at 28 °C, picked and grown o/n in LB at 28 °C. LB media was inoculated with 1:300 o/n *Salmonella* +pKD46 (JVO-3013) culture with 0.2% L arabinose for λRed gene induction and expression. At OD 0.3 cells were placed on ice for 30 min and the culture was pelleted at 4,000 × g for 20 min at 4 °C and washed with ice-chilled water. The cell pellet was dissolved in 1/100 volume of initial culture. 50 μl of competent cells were mixed with 100-300 ng of the PCR product in a chilled 0.2 cm electroporation cuvette and transformed (200 Ω, 25 μF, and 2.5 kV). Transformed cells were recovered in 500 μl prewarmed LB media at 37 °C and incubated for 2 h. Cells were pelleted at 6,000 × g for 3 min at 25 °C and selected by kanamycin resistance on a plate at 37 °C.

Single colonies were verified by PCR amplification of the cassette and chromosomal elements aside by verification primers, and selective growth with kanamycin and not with ampicillin; hence, the pKD46 plasmid was lost. To ensure only single insertion in the chromosomal region, P22 phage lysis and chromosomal integration was performed. The recipient strain (JVS-1574) was P22 phage lysed and the phage lysate was used for lysis of the donor strain. The donor strain phage lysate was then again used for transduction in the recipient strain. In detail, a single colony of the recipient strain was inoculated in 5 ml LB media and grown to OD 1.0. 10 μl of P22 lysate were added and the cells were grown o/n at 37 °C. The suspension was pelleted at 14,000 × g for 10 min at 4 °C. The supernatant was transferred to glass tubes and extracted with 200 μl chloroform to kill all remaining bacteria. The recipient P22 lysate was stored at 4 °C and used for lysis of the donor strain.

For integration of the fragment from the P22 lysate into the recipient strain, 100 μl of OD1 recipient strain were infected with 10-30 μl phage lysate for 20 min at room temperature. The infection was quenched with 20 mM EGTA to allow only one round of infection. The cells were recovered in 1 ml LB media for 2 h and plated on plates with kanamycin. The kanamycin cassette was removed by FRT recombination and transformation of pCP20 at 28 °C. pKD46 and pCP20 were removed by o/n incubation at 42 °C and selection of single colonies on plates with corresponding antibiotics. Strains that were only growing on LB plates were considered as cured and used for studies.

### Western blotting

Protein samples were diluted in reducing 1× protein-sample loading buffer (60 mM Tris-HCl pH 6.8, 0.2 g/ml SDS, 0.1 mg/ml bromphenol blue, 77 mg/ml DTT, 10% glycerol) and boiled at 100 °C for 5 min and loaded on SDS-PAGE. In case of Hfq-FLAG samples were heated to 90 °C. The gel was blotted by semi-dry transfer on polyvinylidene fluoride (PVDF) membranes and probed with tag-specific antibodies. For FLAG-tag detection, a monoclonal anti-body was used (#F1804, 1:2,000 dilution, Sigma Aldrich). GroEL was detected after stripping with anti-GroEL rabbit anti-body (#G6532, 1:10,000 dilution, Sigma Aldrich).

### Co-Immunoprecipitation (Co-IP)

60 OD of cells at OD 2 were resuspended in 800 μl lysis buffer (20 mM Tris/HCl pH 8.0, 150 mM KCl, 1 mM MgCl_2_, 1 mM DTT) and 800 μl 0.1 mm glass beads (BioSpecs). Cells were lysed at 30 Hz for 10 min in the Retsch MM200 at 4 °C. The lysate was cleared twice for 10 min at full-speed at 4 °C. 35 μl monoclonal anti-FLAG M2 antibody produced in mouse (Sigma-Aldrich) was added to the supernatant and incubated for 30 min at 4 °C. 75 μl pre-washed protein A sepharose (Sigma-Aldrich) was added for recovery of FLAG-antibody and incubated for 30 min. Beads were pelleted at 300 × g and washed 5 times witch lysis buffer, resuspended in 500 μl lysis buffer and extracted with PCI. The aqueous layer was precipitated with ethanol and 1:30 3 M sodium acetate at pH 5.2 at −20 °C over-night. The precipitated RNA was pelleted and washed with 500 μl 75% ethanol. The pellet was dried and solubilized in 15.5 μl water. DNA was degraded by DNase I (Thermo Scientific) as described in the manual in presence of RNase inhibitor for 30 min at 37 °C. The sample was diluted with 100 μl water and the RNA was extracted by PCI. The RNA was solubilized to an OD_260_ of 1 and analysed in the Bioanalyzer (Agilent) for quality and quantity estimation for RNA-seq. Protein samples were recovered from the organic layer by 10 vol methanol precipitation.

### RNA-seq

RNA library preparation was conducted by the Core Unit SysMed (Kristina Döring, Würzburg). 200 ng eluted RNA from the coIP were denatured at 70 °C for 2 min, cooled down on ice, and 3’-end dephosphorylated by 10 U T4-PNK (M0201S, NEB) in the presence of 20 U RNase inhibitor (M0314L, NEB) for 60 min at 37 °C. 1.6 mM ATP and 10 U PNK were added for 5’-end phosphorylation for 60 min at 37 °C. RNA was purified by silica-based columns (Zymo RNA Clean & Concentrator kit, R1013, Zymo Research) and eluted in 15 μl nuclease-free water (436912C, VWR). RNA was denatured again and pyrophosphates were removed from the 5’-end by 10 U RppH (M0356S, NEB) in 1× NEB buffer 2 (10 mM Tris-HCl pH 7.9, 50 mM NaCl, 10 mM MgCl_2_, 1 mM DTT) in the presence of 20 U RNase inhibitor for 60 min at 37 °C, and again column purified and eluted in 6 μl. For 3’ adaptor ligation the NEBNext Small RNA Kit (E7560S, NEB) was used: 200 ng RNA were mixed with 1 μl 1:3 diluted 3’ SR adaptors for Illumina, and denatured, 1× 3’ ligation reaction buffer was added and 3 μl ligation enzyme mix in a total volume of 20 μl for 60 min at 25 °C. 1 μl 1:3 diluted SR RT-primer for Illumina were added in a total volume of 25 μl. The annealing was conducted at 75 °C for 5 min, 37 °C at 15 min, 25 °C at 15 min, cooling at 4 °C. 1 μl 5’ SR adaptor for Illumina (freshly solubilized, denatured, and cooled on ice), 1 μl 10× 5’ ligation reaction buffer, and 2.5 μl 5’ ligation enzyme mix were added to the reaction and incubated for 60 min at 25 °C. For reverse transcription, 8 μl first strand synthesis reaction buffer, 1 μl murine RNase inhibitor, and 1 μl ProtoScript II reverse transcriptase were added and incubated for 60 min at 50 °C. PCR amplification of reverse transcripts was conducted by addition of 50 μl LongAmp Taq 2× master mix, 5 μl nuclease free water, 2.5 μl 1:3 diluted SR primer for Illumina, and 2.5 μl 1:3 diluted index primers. The reaction was cycled at 94 °C for 30 s, 14× 94 °C for 15s – 62 °C for 30 s – 70 °C for 90 s, 70 °C for 5 min, and cooled at 4 °C.

DNA was purified by MagSi-NGSPREP Plus (MDKT00010005, Steinbrenner Laborsysteme GmbH) beads with a ratio of 1.8× to reaction volume. All steps were executed in DNA-LoBind tubes (#0030108051, Eppendorf). The final supernatant was analysed by Qubit 2.0 (Thermo Scientific) with the dsDNA HS Assay Kit (Q32854, Thermo Scientific) and the Bioanalyzer 2100 (G2939BA, Agilent) with the DNA 1000 kit (5067-1504) or HS DNA 7500 kit (5067-1506). The sample was sequenced on the NextSeq-500 system (Illumina) with a read depth of 5 mio. reads per sample and a read length of 75 nt single end.

### RNA-seq analysis

Reads were mapped to the *Salmonella* reference sequences (NC_016810.1, NC_017718.1, NC_017719.1, and NC_017720.1) by the READemption *align* function (READemption version 0.4.3, (Förstner et al. 2014)). Coverage wig-files were generated with the *coverage* function and read allocation to genomic features was quantified by the *gene_quanti* function. Annotation of 3’/5’UTRs was used from the Hinton lab and sRNAs from the Vogel lab. Overlaid annotations that caused multiple entries were manually corrected. Sequencing coverages were visualized in the Integrative Genomics Viewer (IGV, Broad Institute, (Robinson et al. 2011)) based on uniquely mapped reads and normalized to the total number of aligned reads.

### RNA preparation

4 OD (for OD2 + 6h, 16 OD of cells were used) of *Salmonella* cells were harvested by mixture with 0.2 volume stop mix (95% (v/v) ethanol, 5% (v/v) phenol, >pH 7.0) and flash freezing in liquid nitrogen. Samples were thawed, pelleted and the supernatant completely removed. The cell pellet was resuspended in 600 μl resuspension buffer (TE pH 8.0, with 0.5 mg/ml lysozyme) and transferred to a 2 ml tube. 1% SDS was added and mixed at 64 °C for 2 min by inversion. 0.3 M NaOAc pH 5.2 was added and mixed by inversion.

Extraction was started by addition of 750 μl acidic phenol and incubation at 64 °C for 6 min with strong shaking. The samples were chilled on ice, and the phases separated by centrifugation at full speed for 15 min at 4 °C. The aqueous layer was transferred and re-extracted with 750 μl chloroform by inversion, and centrifugation. The aqueous layer was precipitated with 2 vol 30:1 mix (ethanol:3 M NaOAc pH 6.5). RNA was precipitated over night at −20 °C and pelleted at full speed for 15 min at 4 °C. The pellet was washed with 70% (v/v) ice cold ethanol, and the resulting pellet was dried at 60 °C. The RNA was resuspended in DEPC treated water at 60 °C, and the quality checked by 1% agarose, 1×TBE gel electrophoresis, and ethidium bromide staining.

### RNA T7-transcription

Template DNA for transcription from a T7 promoter was produced by PCR and extracted from agarose gel or purified by anion-exchange columns, or reverse-complementary primers were annealed as DNA template. RNA was produced with the T7 MEGAscript kit (Thermo Scientific), 200 ng DNA template was mixed in a 20 μl transcription reaction, with 2 μl of each nucleotide, 2 μl enzyme mix, and 2 μl 10× reaction buffer, and incubated for 6 h to over-night at 37 °C. 1 μl Turbo DNase I (1 U/μl) was added and incubated for 15 min. 15 μl 5 M ammonium acetate, 100 mM EDTA (pH 8.0) were added and the sample was filled up to 100 μl with water and extracted with PCI. The aqueous phase was rebuffered through a G25 spin-column and sample was eluted in water, and precipitated with 7.5 mM NaOAc pH 5.2 and 75% ethanol. The pellet was washed with 75% ethanol, air-dried, and solubilized in nuclease-free water. RNA quantity and quality was assayed by Urea-PAGE and Stain-All (0.075 mg/ml in 60% formamide, autoclaved, E9379, Sigma-Aldrich) staining.

### 5’ ^32^P-labelling of RNA

RNA for EMSA, structure probing, or kinetic studies, or DNA oligo for northern probing were radioactively labelled by polynucleotide kinase. In a total reaction of 10 μl, up to 100 pmol RNA or DNA were mixed with 2.5 pmol γ-^32^P-ATP (1 μl, 400 mCi/μmol, 10 μCi/μl, Hartmann Analytic) in 1× kinase buffer (80 mM Tris-HCl pH 7.5, 10 mM MgCl_2_, 5 mM DTT), and 1 μl T4-PNK (10 U/μl, EK0031, Thermo Scientific). The reaction was incubated at 37 °C for 40 min. Free nucleotides were removed with a spin column (Illustra MicroSpin G-25 Columns, GE Healthcare) at 750 × g for 2 min. The labelled RNA/DNA was eluted in 20 μl water. Alternatively, the reaction was extracted by PCI for nucleotides (A156.3, Carl Roth), and precipitated by ethanol.

### FopA protein expression and purification

A _H6-3C_FopA expression plasmid (pMiG-006, pET-M14+ backbone, kanamycin resistance, cloned by ‘Recombinant Protein Expression Facility’, Rudolf-Virchow-Center) was transformed into *E. coli* BL21-CodonPlus (DE3)-RIL cells (Agilent Technologies, JVS-12280, chloramphenicol resistance). Cells were grown in LB to OD_600_ 0.6 at 37 °C, cooled to 18 °C and induced with 0.5 mM IPTG over-night. Cells were resuspended in lysis buffer (50 mM sodium phosphate pH 8.0, 1 M NaCl, 1 mM TCEP, 1 mM PMSF) and disrupted by sonication (50% amplitude, 30 s pulsation - 30 s break for 5 min, on ice, Sonopuls HD 3200, TT13 tip, Bandelin). The lysate was cleared at 15,000 × g for 15 min at 4 °C. Equilibrated Protino Ni-IDA beads (Macherey-Nagel) were added to the supernatant and incubated for 1 h at 4 °C in an over-head rotor. Beads were recovered and washed with lysis buffer.

Protein was recovered in elution buffer (50 mM sodium phosphate pH 8.0, 200 mM imidazole, 1 mM TCEP), rebuffered by centrifugal filtration (Amicon Ultra, Merck Millipore, cut-off 10 kDa) in 3C-cleavage buffer (50 mM Tris/HCl pH 7.5, 150 mM NaCl, 0.1 mM EDTA, 1 mM TCEP) and incubated over-night with 3C protease at 4 °C. Ni-IDA beads were added to the sample for removal of the cleaved tag and the protease. The supernatant was diluted 1:50 in cation exchange buffer A (50 mM sodium phosphate pH 6.0, 20 mM NaCl, 0.1 mM EDTA, 1 mM TCEP). The sample was applied to a mono S 5/50 GL (GE) column and eluted in a salt gradient to 1 M NaCl. FopA eluted at about 0.7 M NaCl. The protein was concentrated and rebuffered in storage buffer (20 mM sodium phosphate pH 7.0, 100 mM NaCl) through centrifugal filtration and flash frozen in liquid nitrogen. The concentration of FopA was determined spectrophotometrically utilizing the extinction coefficient of 21,000 mM^−1^cm^−1^.

### Electrophoretic mobility shift assay

4-8 nM 5’-^32^P-labeled RNAs (indicated by a *) were mixed optionally with equal amount of *repZ* and incubated for 5 min at 37 °C in reaction buffer (25 mM Tris/HCl pH 7.4, 150 mM NaCl, 1 mM MgCl_2_). 0.1 mg/ml yeast RNA (10 mg/ml, Merck) was added. Final concentration of 0-5 μM FopA was added and incubated for 15 min at 37 °C. Subsequently, 3 μl 5× native loading dye (0.5× TBE, 50% (v/v) glycerol, 0.2% (w/v) xylene cyanol, 0.2% (w/v) bromphenol blue) were added and loaded immediately on a running native 6% polyacrylamide/0.5× TBE gel and run for 2-3 h at 40 mA at 4 °C. The gel was dried on filter-paper. The membrane was wrapped in foil and exposed to a photo-stimulatable phosphor plate and the signal was detected by the Typhoon 7000 phosphoimager (GE Healthcare).

### RNA-RNA complex formation kinetics

Kinetics of complex formation between Inc and *repZ* were determined as described previously (Persson et al. 1988; Asano et al. 1998). 1 nM ^32^P-labeled Inc RNA (denatured by boiling and cooled to 37 °C) was mixed optionally with 200 nM FopA and incubated for 10 min at 37 °C in reaction buffer (20 mM Tris/HCl pH 7.5, 1 mM MgCl_2_, 150 mM NaCl). 25 nM *repZ* was added and immediately time points were taken by dilution with 2 volumes in application buffer (92% deionized formamide, 17 mM EDTA, 0.025% xylene cyanol, 0.025% bromphenol blue) and loading onto a running and cooled 8.3 M Urea-PAGE gel at 40 W for 2 h. For the 0 s time point, Inc(±FopA) was quenched with 2 volumes application buffer and then *repZ* was added and loaded immediately onto the running gel. The kinetics of the complex formation were calculated by the pseudo-first-order rate constant *k*_1_’ = ln(2)/t_1/2_ (t_1/2_ was determined from the plot) that is related to the second-order rate constant *k*_2_ = *k*_1_’/[*repZ*], as described previously (Persson et al. 1988).

### Structure probing

5’-^32^P-labeled Inc (Inc*) and *repZ* RNA were denatured at 95 °C for 3 min and cooled slowly down to 37 °C in reaction buffer (25 mM Tris-HCl pH 7.4, 150 mM NaCl, 1 mM MgCl_2_). 15 nM Inc* was incubated optionally with 15 nM *repZ* at 37 °C for 5 min in reaction buffer. A final concentration of 0.1 mg/ml yeast RNA (AM7118, Thermo Scientific) was added, incubated for 1 min, and optionally 5 μM of FopA (His-tag cleaved off by 3C protease) were added to a final volume of 10 μl and incubated for 15 min at 37 °C to allow binding. In a total reaction of 12 μl, 0.002 U/μl RNase T1 (1 U/μl, AM2283, Thermo Scientific) was added and incubated for 3 min, or 0.2 ng/μl RNase A (10 mg/ml, EN0531, Thermo Scientific) for 3 min. For the ladder, 30 nM Inc* RNA was incubated for 5 min at 95 °C in 10 μl alkaline hydrolysis buffer (50 mM sodium carbonate pH 9.2, 1 mM EDTA), or for 3 min at 37 °C with 0.01 U/μl RNase T1. The reactions were quenched with 50 mM EDTA, 0.3 M NaOAc pH 5.2, 50% formamide (deionized, P040.1, Carl Roth), and 2% SDS. The samples were boiled for 5 min at 95 °C, cooled down on ice. Samples were diluted with water and extracted with an equal volume of PCI for RNA (X985.3, Carl Roth). RNA in the aqueous phase was precipitated over-night at −20 °C with 80% ice-cold ethanol, 0.06 M NaOAc pH 5.2, 0.2 mM MgCl_2_, and 1 μl GlycoBlue (15 mg/ml, AM9515, Thermo Scientific) was added as tracer. After pelleting at full-speed and 4 °C, the pellet was washed with ice-cold 75% ethanol, and the supernatant removed, and evaporated. The RNA was solubilized in GLII buffer (95% deionized formamide, 0.02% SDS, 0.02% bromphenol blue, 0.01% xylene cyanol, and 1 mM EDTA) for 5 min at 60 °C, boiled at 95 °C for 5 min. The hot sample was loaded onto a pre-run (45 W for 1 h) and hot 10% 7 M Urea-PAGE gel in 1× TBE buffer (89 mM Tris, 89 boric acid, 2 mM EDTA, pH ~8.3). When the running front reached nearly the end. The gel was transferred onto filter-paper, sealed from the top with foil, and dried in the Gel Dryer Model 543 (Bio-Rad) for 1 h with heating. The gel was set onto a photostimulatable storage phosphor screen and the signal was detected by the Typhoon 7000 phosphorimager (GE Healthcare).

## DATA AVAILABILITY

MS data are accessible at the ProteomeXchange consortium (Deutsch et al. 2016) via the PRIDE partner repository (Perez-Riverol et al. 2018) with the dataset identifier PXD018422. Reviewers can access the dataset at https://www.ebi.ac.uk/pride/archive with the login details user: and the password: ‘cXGcSEoO’. Raw data after MaxQuant analysis are listed in Supplemental Table 1 in the tab ‘RAW’.

The python code for the GradR browser is deposited at Zenodo 3742229 (DOI:10.5281/zenodo.3742229) and accessible for review process under the following shared link without registration: https://zenodo.org/record/3742229?token=eyJhbGciOiJIUzUxMiIsImV4cCI6MTYxNzIyNzk5OSwiaWF0IjoxNTg2MjUxMzg1fQ.eyJkYXRhIjp7InJlY2lkIjozNzQyMjI5fSwiaWQiOjcxNDEsInJuZCI6Ijc4ZjUxYTg1In0.YVD9bR01KUVZk7oQsWHPC14xH1ozruC9LrPJqyQQQkUIXVK1HliiohiE7sy9rco72Io9oSql3AAm5m-SPVGjwg

The GradR browser is online accessible at www.helmholtz-hiri.de/en/datasets/GradRSeT with the login details user: ‘reviewer’ password: ‘welcome’. FopA coIP-seq raw FASTQ and analysed WIG coverage files are accessible at Gene Expression Omnibus (GEO, (Edgar et al. 2002)) with the accession no. GSE148184. Reviewers can enter the dataset with the secure token ‘sdmbegqylrkfloz’.

READemption 0.4.5 is deposited at Zenodo 1134354 (DOI:10.5281/zenodo.1134354). The READemption analysis folder is deposited at Zenodo 3742725 (DOI:10.5281/zenodo.3742725) and accessible for review process under the following shared link without registration: https://zenodo.org/record/3742725?token=eyJhbGciOiJIUzUxMiIsImV4cCI6MTYxNzIyNzk5OSwiaWF0IjoxNTg2MjY0MTYzfQ.eyJkYXRhIip7InJlY2lkIjozNzQyNzI1fSwiaWQiOjcxNDYsInJuZCI6IjgyNGNhYWU4In0.Pp6xsXIREjePiJuqLBVy1SYB1MzIrZy59GvRW8R-7SIcYJ0ExxD7wIpoX-TzANHzKuLmNPebKF71V4mWbpWIFw

## Supplemental material

**Supplemental Fig. S1.**
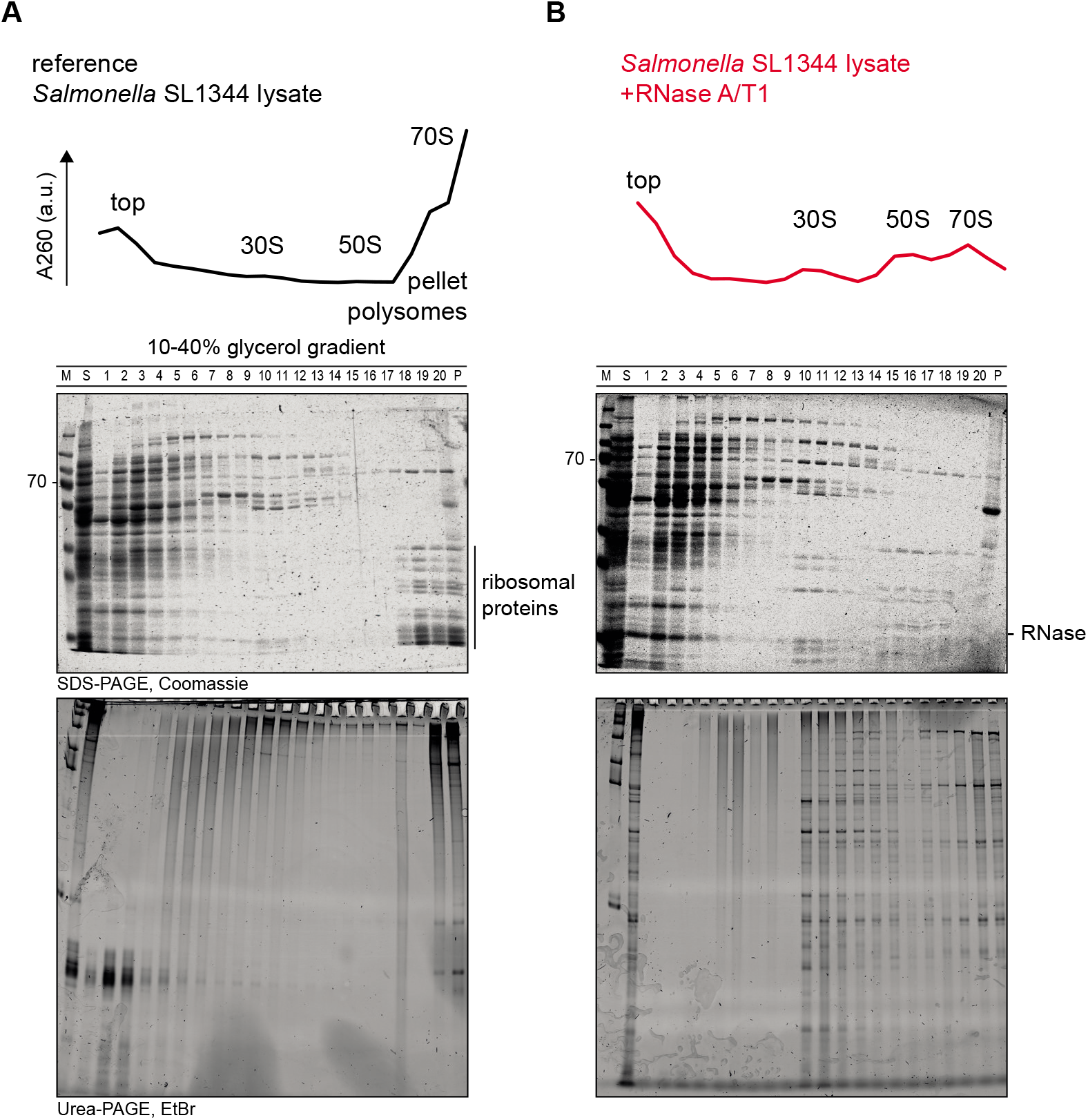
GradR protein and RNA distribution. (A), (B), A_260_ profile, protein and RNA patterns in reference and +RNase gradients were resolved in SDS-PAGE and 6% Urea-PAGE.

**Supplemental Fig. S2.**
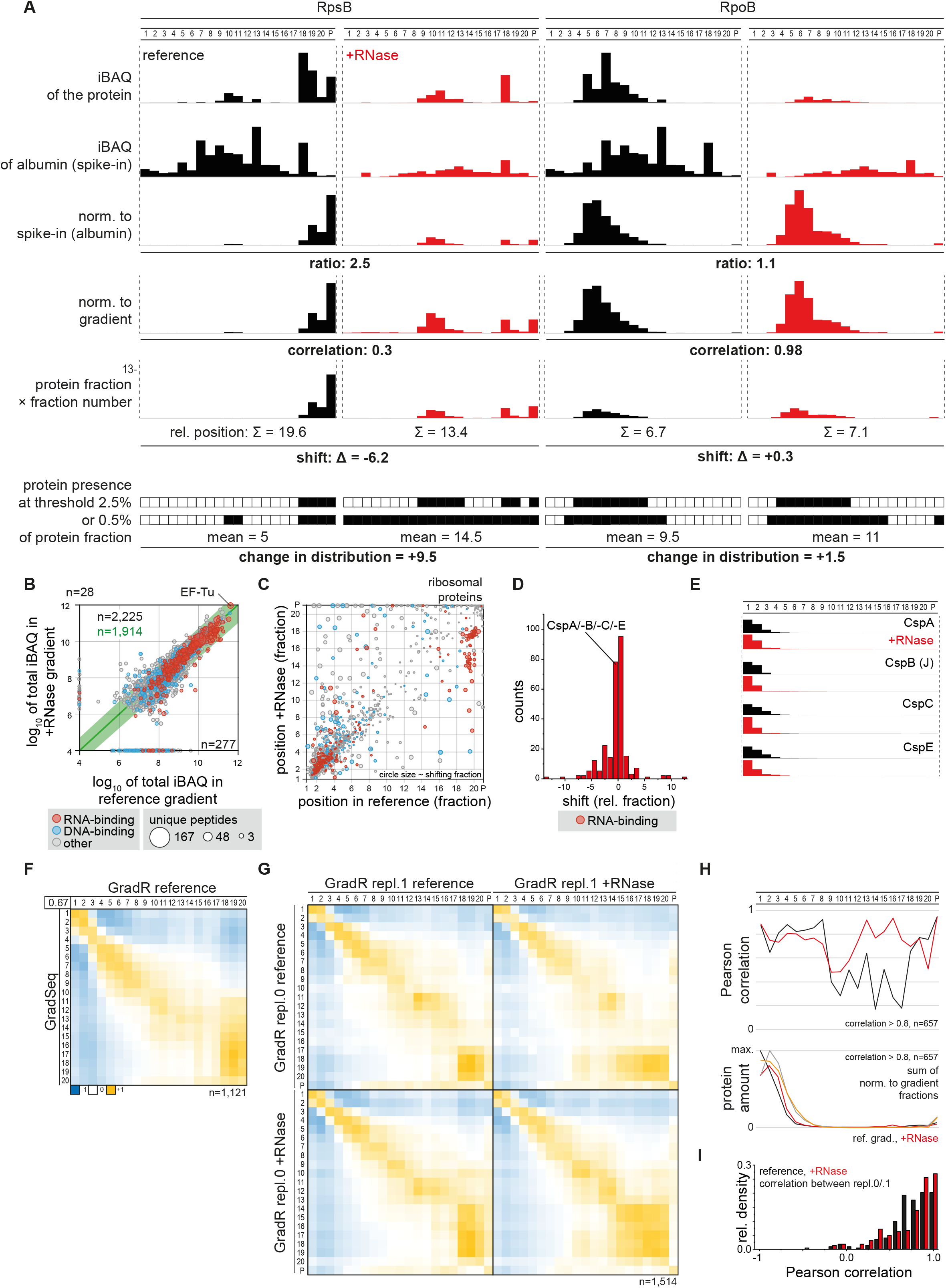
GradR protein and RNA distribution. (A) Protein abundance normalization and calculation of shift and distribution changes for RpsB (30S) and RpoB (RNAP). (B) Total protein levels in reference and +RNase gradients were similar (5-fold difference in abundance is in the green boundary) (C) Relative position calculated for both gradients dependent on the protein amount in each fraction. Ribosomal proteins accumulate in HMW fractions. Ribosomal small and large subunits can be discriminated by a shift of 4 or 2 relative fractions, respectively. The majority of the proteins allocated in the first gradient half. (D) Not all proteins that are classified as *RNA-binding* (red) are shifting, a majority is allocating around 0 shifting relative fractions. (E) For example, the cold-shock proteins are shifting only marginally. (F) Correlation of reference gradient with Grad-seq. Ribosome dissociate in Grad-seq, hence the correlation is diminished in ribosomal subunit fractions. (G) During development of GradR, we performed an initial experimental replicate 0 with modified conditions: the lysis buffer contained 0.2% Triton X100 as in Grad-seq (Smirnov et al. 2016). The protein samples were prepared for MS detection by acetone precipitation and solubilized directly in digestion buffer with 8 M urea. All other steps were performed as stated before for replicate 1. GradR replicates with 0.2% Triton X100 (repl. 0) and without (repl. 1) correlated well. (H) Fraction-wise correlation coefficient in (G) was about 0.8 in fractions with high protein load. (I) Distribution of correlation coefficients of sedimentation profiles across replicate experiments in (G,H) shows that the majority of (non-shifting) proteins were highly correlated.

**Supplemental Fig. S3.**
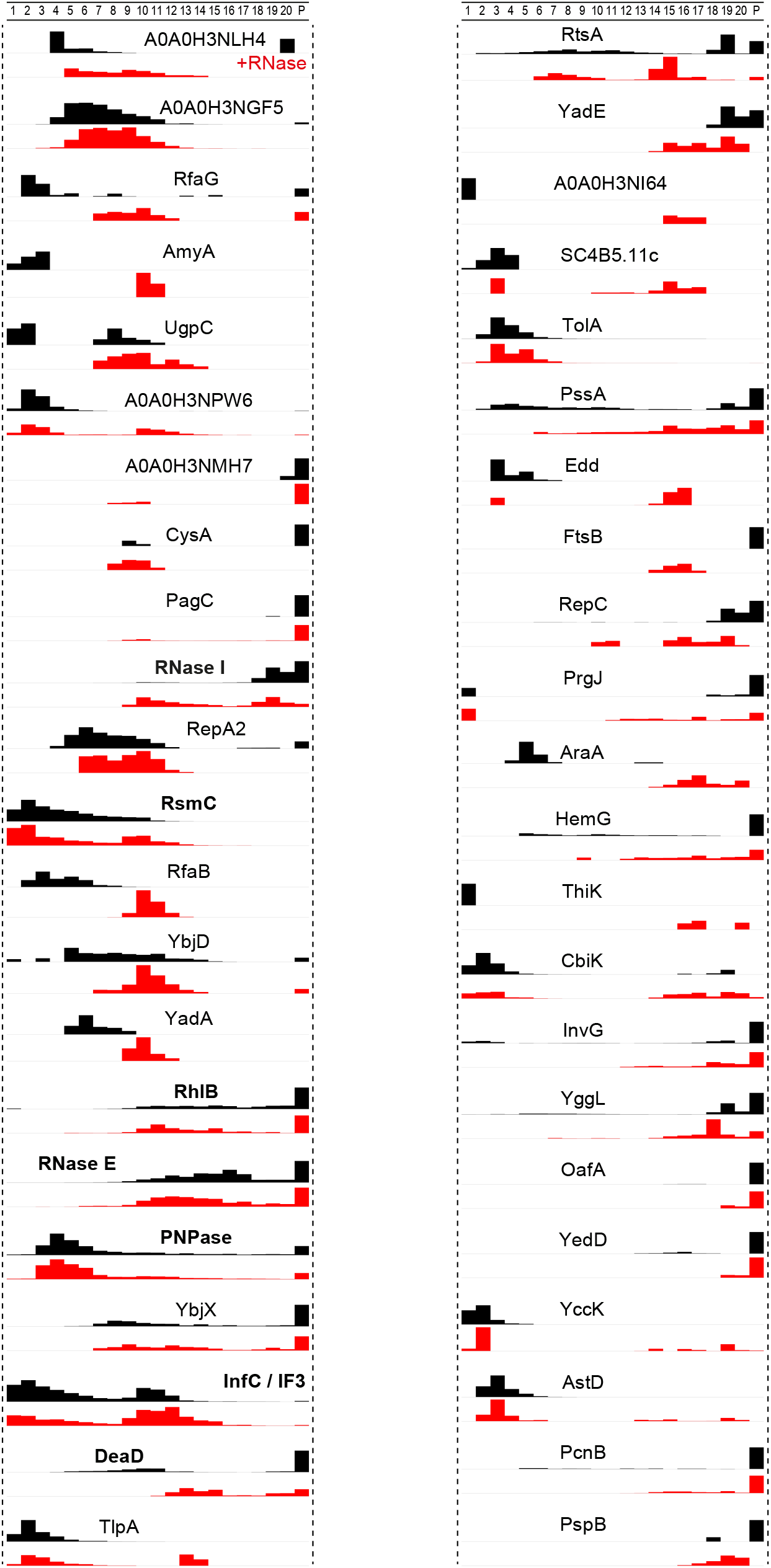
Ribosome associated proteins enrich at ribosomal fractions after RNase digestion. Proteins that shifted towards fractions 10 to 20 are represented. Transiently associated ribosomal protein are indicated in bold type. Other proteins represent novel ribosome-associated candidates, among them six are fully uncharacterized and named by UniProt-ID.

**Supplemental Fig. S4.**
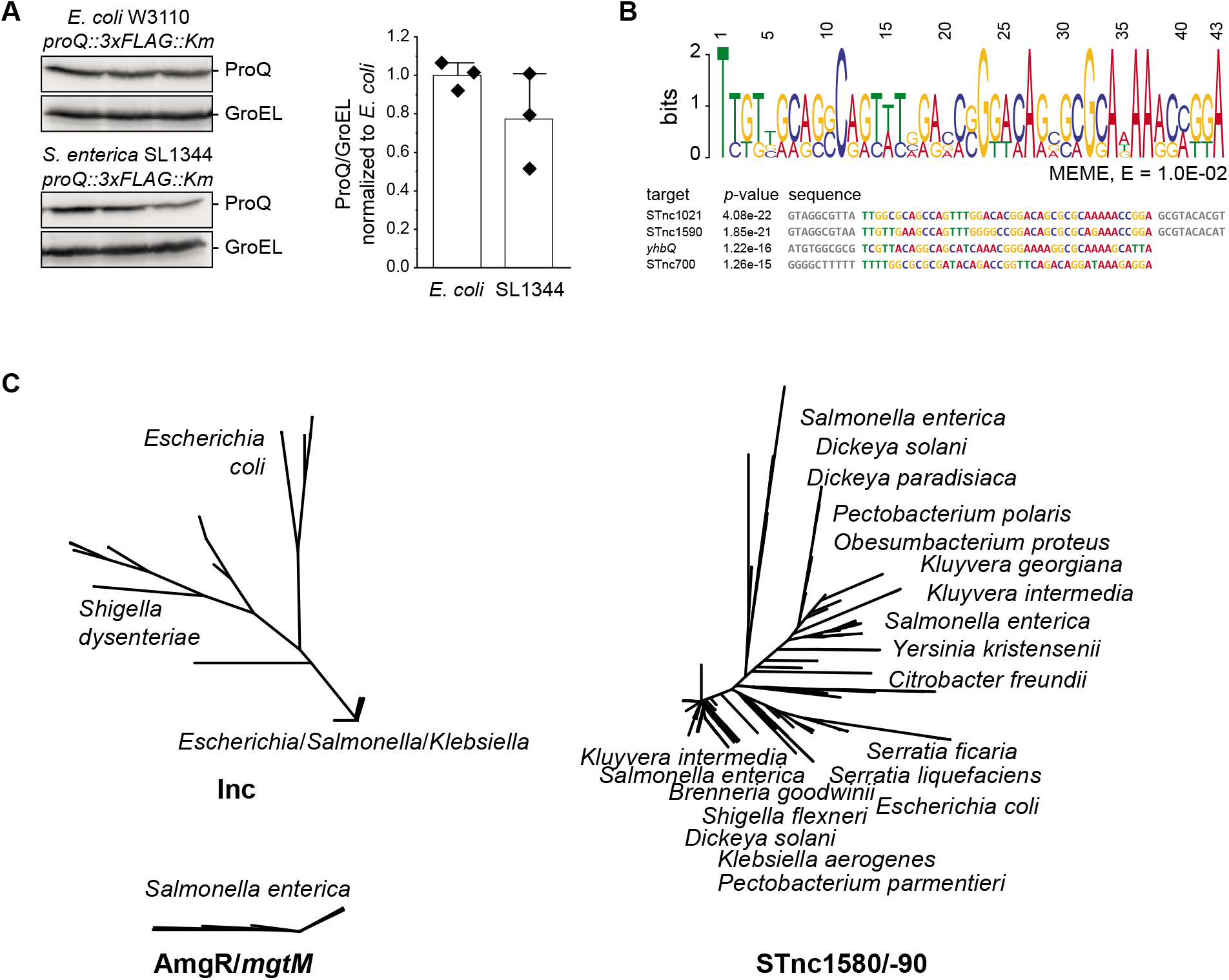
ProQ protein levels. (A) ProQ levels in *E. coli* and *S. enterica* were in the same range. (B) Motif-based sequence analysis (MEME) did not yield a significant sequence motif that would match the top recovered targets in Fig. 5B,C. (C) Read-covered sequences of precipitated RNAs by FopA were submitted to GLASSgo (Lott et al. 2018) for RNA homolog search. The resulting sequences were aligned for calculation of phylogenetic distances by Clustal Omega (EMBL) and visualized with FigTree (1.4.4, Rambaut A, University of Edinburgh).

**Supplemental Fig. S5.**
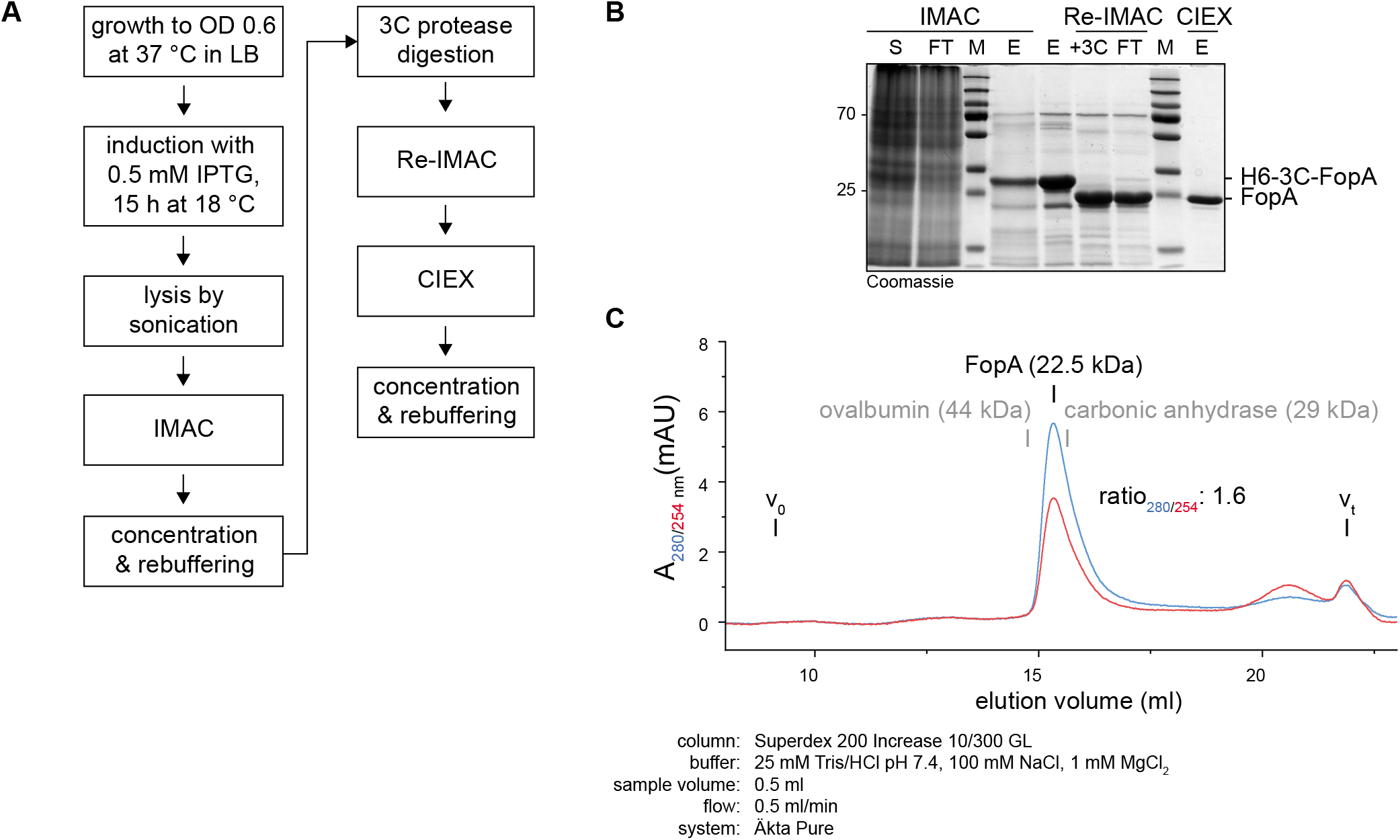
FopA purification. (A) General purification procedure for FopA. (B), Immobilized metal-affinity chromatography (IMAC) captured His-tagged _H6-3C-_FopA. The N-terminal His-tag was cleaved off by 3C protease and removed by reverse-IMAC. In a cation-exchange chromatography (CIEX) at pH 6.0, FopA eluted in RNA-free form at 0.6 M NaCl. (C) In size-exclusion chromatography, FopA eluted as a monomer with a 280/254 nm absorption ratio of 1.6.

**Supplemental Fig. S6.**
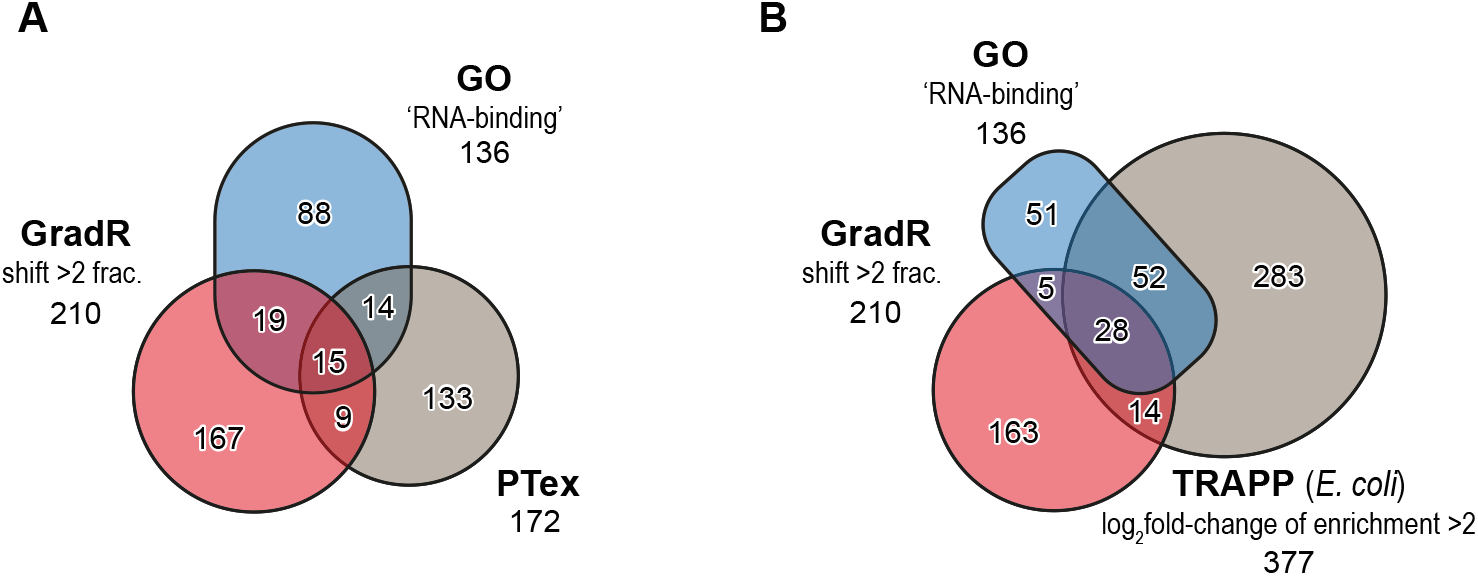
GradR is an orthogonal RBP discovery method to UV-crosslinking based approaches. (A) Phenol-Toluol-based extraction of RPBs (PTex, (Urdaneta et al. 2019)) predicted 24 shared candidates of which 15 were GO-term annotated as RBPs. (B) Silica-based recovery of RNA-RBP candidates (TRAPP, (Shchepachev et al. 2019)) yielded 42 matching candidates of which 28 were GO-term annotated as RBPs. The correlations between the here listed methods were only minor and indicate orthogonality. Candidate numbers are based on applied thresholds in the individual studies. *Salmonella* and *E. coli* RBP candidates from GradR and TRAPP were matched by gene name if applicable.

**Supplemental Fig. S7.**
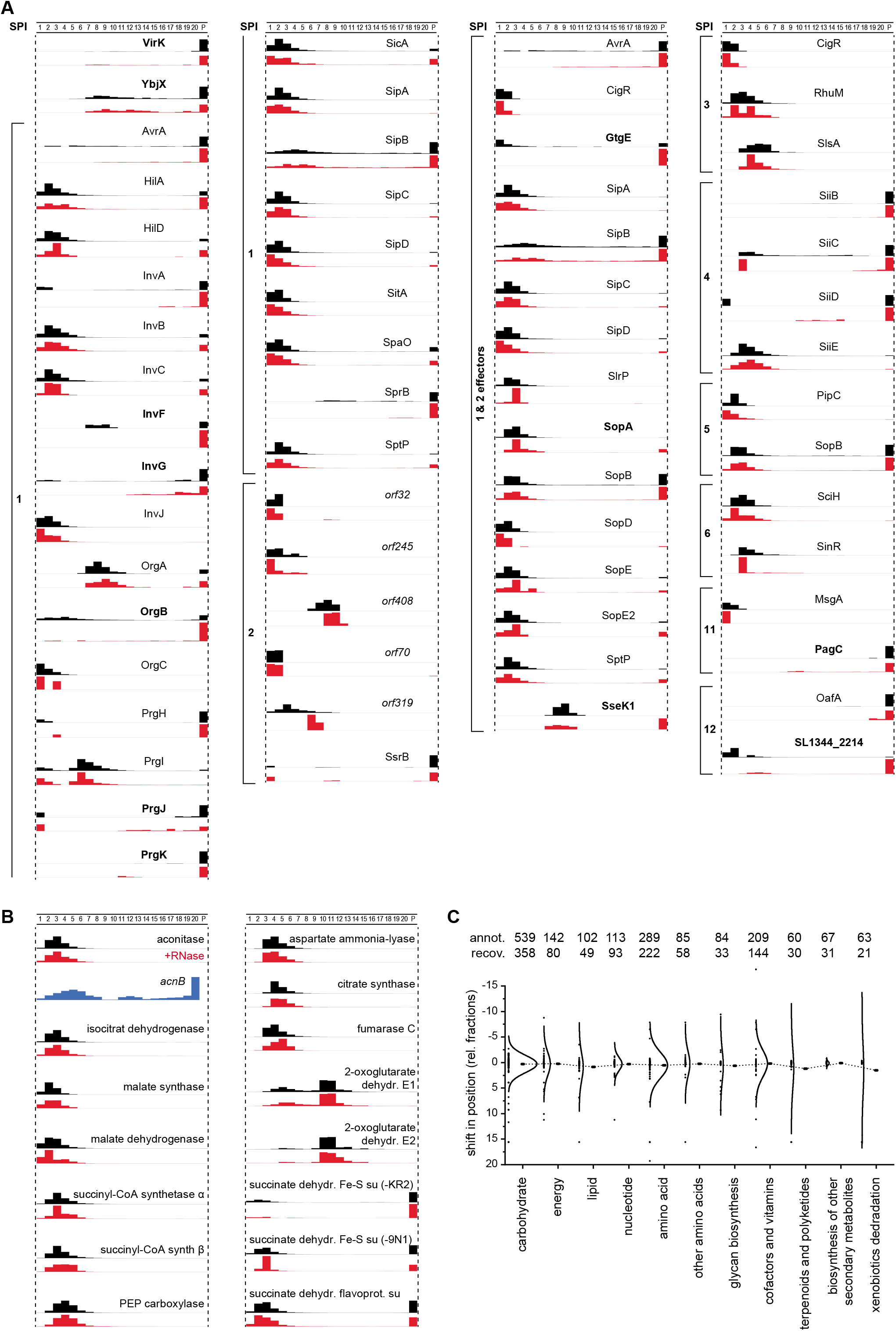
A number of virulence factors shifted whereas metabolic protein sedimentation was not affected by RNase treatment. (A) Virulence factors were grouped as encoded in *Salmonella* pathogenicity islands (SPI). A number of factors that shifted after RNase treatment are represented in bold. (B) Aconitase and enzymes from the TCA cycle were not shifting upon RNase treatment. (C) A global analysis of metabolic proteins based on KEGG categories yielded no general shifting behaviour.

**Supplemental Table S1 I List of sedimentation profiles for detected proteins.**

Supplemental_TableS1_GradR_data_with_analysis_pipeline.xlsx

**Supplemental Table S2 I coIP-seq results of FopA at OD2.**

Supplemental_TableS2_FopA_coIP-seq.xlsx

**Supplemental Table S3 I STAR Methods: oligonucleotides, plasmids, and strains used in the study.**

Supplemental_TableS3_STAR_methods.xlsx

**Supplemental Table S4 I Classifications used in this study.**

Supplemental_TableS4_classficiation_lists.xlsx

